# Mixed-model admixture mapping identifies smoking-dependent loci of lung function in African Americans

**DOI:** 10.1101/626077

**Authors:** Andrey Ziyatdinov, Margaret M. Parker, Amaury Vaysse, Terri H. Beaty, Peter Kraft, Michael H. Cho, Hugues Aschard

## Abstract

Admixture mapping has led to the discovery of many genes associated with differential disease risk by ancestry, highlighting the importance of ancestry-based approaches to association studies. However, the potential of admixture mapping in deciphering the interplay between genes and environment exposures has been seldom explored. Here, we performed a genome-wide screening of local ancestry-smoking interactions for five spirometric lung function phenotypes in 3,300 African Americans from the COPDGene study. To account for population structure and outcome heterogeneity across exposure groups, we developed a multi-component linear mixed model for mapping gene-environment interactions, and empirically showed its robustness and increased power. When applied to the COPDGene study, our approach identified two 11p15.2-3 and 2q37 loci, exhibiting local ancestry-smoking interactions at genome-wide significant level, that would have been missed by standard singlenucleotide polymorphism analyses. These two loci harbor the *PARVA* and *RAB17* genes previously recognized to be involved in smoking behavior. Overall, our study provides the first evidence for potential synergistic effects between African ancestry and smoking on pulmonary function and underlines the importance of ethnic diversity in genetic studies.

## Introduction

The study of genetically diverse populations has become a priority in public health research. Several major initiatives started in the past few years, including the National Institute of Health (NIH) Trans-Omics in Precision Medicine (TOPMed) Program that aims to sequence over a hundred of thousands whole genomes from a variety of ancestries. These initiatives compensate for the lack of participants of non-European ancestries in genetic studies^1–3^. Besides addressing health disparities among ethnic groups, studies of multi-ethnic cohorts and admixed populations can provide important information about the biology of complex diseases and help to identify associated genes^4^. Recently admixed populations, such as African Americans, represent a special case of multi-ethnic cohorts with mosaic chromosomes derived from several ancestral populations. Admixture mapping, often applied to recently admixed populations, searches for genomic loci of unusual local ancestry at a putative disease risk locus compared with the genome-wide average^5^. Findings for respiratory disease, chronic renal disease, prostate cancer and systemic lupus erythematosus have been reported as results of admixture mapping^7–12^.

The data analysis in admixture mapping consists of two main steps, inferring local ancestry and testing for association between every local ancestry segment and an observed phenotype. Given unobservable ancestry information, current methods on ancestry inference probabilistically define the location of every ancestral switch using genotyping array data, reference haplotypes and algorithms based on hidden Markov models^6,35^. These methods were empirically shown to produce reliable results on African Americans^40^, as they represent a relatively simple two-way admixture and are well modeled by available reference panels^41^. The standard approach for association testing is similar to genome-wide association studies (GWAS) on single-nucleotide polymorphisms (SNPs) and runs linear regression to estimate the correlation between local ancestry and phenotype. To avoid confounding due to population structure that is inherently present in admixed individuals, most studies also included global ancestry components (i.e. the genome-wide proportions of ancestry derived from each ancestral population) as covariates. Going beyond linear regression, the framework of linear mixed models was applied to quantify individual similarities by ancestry, showing how the phenotypic variance is explained by local ancestry^42^ and linking it to the heritability of complex traits estimated from SNP data^20^. Notably, linear mixed models have not yet been applied in admixture mapping despite several potential advantages^35^.

Recently, we found that the correlation between local ancestry and untyped causal variants can be leveraged to detect distant gene-gene interactions in admixed populations through local ancestry-local ancestry screening^18^. That work also demonstrated that the power of such admixture mapping increases with the number of causal variants within local ancestry tested and with the degree of differentiation of variants between the ancestral populations. Here, we suggest that the same principle can be applied to search for gene-environment interactions. Regarding previous studies of gene-environment interactions, most works focused on interactions between the global ancestry and environmental factors using linear regression^16,19–21^. Since here we sought to screen for local ancestry-environment interactions, a type of admixture mapping seldom explored, we argue that such application might face multiple methodological challenges due to both population stratification and outcome heterogeneity among individual groups stratified by environmental exposure. Hence, the use of linear mixed models will be particularly relevant to account for complex genetic and environmental relationships in admixed individuals.

Application of admixture mapping to lung function phenotypes in African Americans is especially relevant^13^. European Americans and African Americans are well known to show differences in spirometric measures of lung function such as forced expiratory volume in one second (FEV_1_) and forced vital capacity (FVC) with higher values for these two traits for European Americans. Factors partially responsible for these differences include body habitus, early-life development conditions, socioeconomic status and other environmental factors^14,15^. Previous studies showed strong evidence that the proportion of African global ancestry is associated with lower lung function for a given ranges of height and age^13,16^. Also, the higher proportion of African ancestry in African Americans was associated with an additional decrease in lung function for smokers^17^.

To address the aforementioned methodological challenges, we used real data and proceed in a step-wise assessment of multi-component linear mixed model in order to define a robust interaction test of association. More precisely, we conducted a genome-wide scan of local ancestry-smoking interactions for five spirometric lung function phenotypes available in 3,300 African-Americans from the COPDGene study^22^. The search for gene-environment interactions related to pulmonary function phenotypes in the COPDGene study is particularly relevant, as it is one of the largest studies of African-American smokers. In result, our application of the proposed linear mixed model identified two genome-wide significant and five suggestive loci that would have been missed in standard single SNP-based approaches.

## Materials and Methods

### The COPDGene dataset

In the analysis of COPDGene study, we focused on five correlated quantitative pulmonary phenotypes: forced expiratory volume in one second (FEV_1_); forced expiratory volume in one second as a percent of predicted (FEV_1_ % predicted); forced vital capacity (FVC); forced vital capacity as a percent of predicted (FVC % predicted); and the ratio of forced expiratory volume in one second to forced vital capacity (FEV_1_/FVC). We derived two binary smoking exposures from the number of cigarettes smoked per day for gene-environment interactions: current smoker exposure (current smokers vs. former smokers) and heavy smoker exposure (current heavy smokers vs. current moderate smokers). We defined moderate current smokers with 1-14 cigarettes per day on average, while heavy current smokers were defined as with >14 cigarettes per day on average. When using heavy smoker exposure in the analysis, we excluded all subjects who were former smokers.

Locus-specific ancestry or local ancestry was inferred from genotype data as previously described^22^. Briefly, we used the LAMP-LD program^37^ to estimates local ancestry per individual using a hidden Markov model (HMM) algorithm comparing observed genotypes and haplotypes from reference ancestral populations. We parametrized the algorithm with 15 HMM states, a window size of 50 SNPs, and used 99 CEU and 108 YRI unrelated individuals from the 1,000 Genomes Project (Phase III)^38^ as reference panels. Per SNP estimates of African ancestry were further used in two post-processing steps. First, we averaged the local ancestry per individual to derive the genome-wide proportion of African ancestry (i.e. the global ancestry). Second, we estimated local ancestry segments by merging neighboring SNPs with identical values. We next filtered out short ancestry segments of length less than 10,000 bases to mitigate possible artifacts of the inference procedure that might affect further admixture mapping analysis^39^.

### Step-wise model selection for admixture mapping

Consider a quantitative trait stored in a *n*-dimensional vector *y* and a *n×m* data matrix *Z* of local ancestry segments, where *n* is the number of individuals and *m* is the number of local ancestry segments. Let *z_l_* be a column of matrix *Z* corresponding to a single local ancestry segment, and *x_e_* being a *n*-dimensional column vector of a binary environmental exposure. We aim at testing the statistical interaction between the local ancestry segments *z_l_* and the exposure *x_e_* on the phenotype *y*. As discussed in Supplementary Material, we are interested to assess a fixed effect of interaction *z_l_* × *x_e_* using the standard *Wald* test, while controlling for variance related to local ancestry, global ancestry and exposure that might confound the estimate of effect. However, in regards to potentially non-trivial structure in the data, due to both ancestry admixture and potential heterogeneity of outcome across exposure groups, we defined our association model using a step-wise approach, where the model complexity was incremental until reaching the desired criteria of validity. Following standard practices^44^, these criteria were designed 1) to reach a genomic inflation parameter (*λ* close to one, and 2); to achieve an overall shape of the standard quantile-quantile plot (QQ-plot) of the −log_10_(P) matching the expected uniform distribution of p-values for the majority of the tests.

In practice, before evaluating interaction effects, we first assessed the robustness of a standard linear mixed model (LMM) when testing for the marginal effect of local ancestry *z_l_* only:

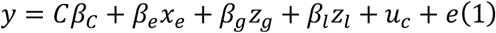

where *C* is a matrix of trait-specific covariates and *β_c_* is a vector of their fixed effects; *z_g_* is a vector of global; and *β_e_, β_g_, β_l_* are fixed effects of exposure, global ancestry and local ancestry, respectively. The random effects include a vector of the residual errors *e* and a vector of random effect *u_c_* encoding whether two given individuals belong to the same medical center. We further added additional random effect components (described below) in the LMM, which importance was assessed incrementally (**Supplementary Table S1**). We next included our parameter of interest, the local ancestry-exposure interaction effect, on top of random effect components selected at the previous step and continued our assessment of additional components until reaching the desired characteristics. Our full and final LMM was defined as follows:

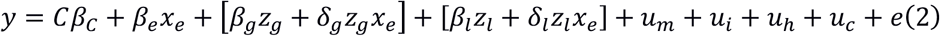

where, in addition to notation in Equation (1), *z_g_* × *x_e_* and *z_l_ × x_e_* represent interactions between global and local ancestries and exposure, respectively; *δ_g_* and *δ_l_* are fixed effects of interactions between global ancestry and local ancestry and exposure, respectively. The first vector of additional random effects *u_m_* captures the variance of local ancestry remaining after taking into account the global ancestry as a fixed effect (*z_g_*)^42,43^. The variance-covariance matrix of *u_m_* is the ancestral relationship matrix (ARM) derived by the cross-product operation on column-wise centered and scaled *Z* matrix^43^. The second vector *u_m_* complements the previous vector *u_m_* and comes out due to testing the interaction effect (*δ_l_*) rather than the marginal effect (*β_l_*)^34^. The variance-covariance matrix of *u_i_* is derived from ARM based on stratification by binary environmental exposure status (*x_e_*)^34^. We refer to this matrix as environmental ARM or EARM. The third vector *u_h_* models the heterogeneity of phenotypic variance across the three smoking groups. The variance-covariance matrix of *u_h_* is a diagonal matrix, where entries are the same if they correspond to the same group.

### Multi-phenotype analysis

We first conducted single-trait admixture mapping of gene-environment interactions for each of the five pulmonary traits considered and each of the two exposures (current smoker and current heavy smoker). However, these five spirometric lung function measurements are all highly correlated and likely share genetic association signals. To reduce the penalty for multiple testing and potentially increase power, we focused our primary screening on a multivariate association tests combining single-trait signals separately for each exposure without assuming any prior on the direction of the single-trait effects^45–48^. In practice, given five Z-scores for a local ancestry segment, stored in a vector*S*_5×B_, we estimated the multivariate Z-score *S_joint_* in two steps. We first estimated *Σ*_5×5_, the variance-covariance matrix among Z-scores under the null hypothesis of no association and then, for each local ancestry segment, we derived the multivariate statistics using the *Mahalanobis* distance, defined as:

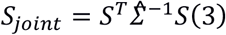

which squared statistics 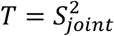 follows a *χ*^2^ distribution with 5 degrees of freedom under the null composite hypothesis of no interaction effect on any of these five traits. For the estimation of *Σ*, previous works suggested using the complete Z-score data from the genome-wide scan, i.e. 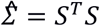, assuming that the vast majority of Z-scores are distributed under the null^46,48^. Here, we made the same assumption and additionally discarded large single-trait Z-scores above a given threshold to reduce the risk of bias. Further, we approximated 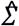 from the resulting truncated multivariate normal distribution by the maximum likelihood estimation. We empirically found the best threshold value equals 3 for our admixture mapping Z-scores (**Supplementary Material**).

### Follow-up analysis

We performed association and further fine-mapping analysis using genotyped data, following up regions identified by admixture mapping. Considering *x_g_* is a vector of genotypes for a single SNP, we extended the interaction model (Equation 2) by adding two terms for marginal genetic (*x_g_*) and interaction (*x_g_ × x_e_*) effects. Note that the exposure term (*x_e_*) was already included in this model. We performed a univariate *Wald* test with one degree of freedom to derive the p-value for interaction effect between genotype and exposure. By including a local ancestry term when testing for the interaction effect of genotype, we accounted for possible different LD patterns for European and African ancestral backgrounds^49^. Such conditional analysis can reduce power but assures that the interaction effect of genotype is driven by a biological mechanism rather than a better SNP tagging in a particular ancestral population.

As discussed in our previous work^18^, we expected SNPs in regions of local ancestry-smoking interactions to show multiple-SNP effects on the trait as well as high allelic frequency differentiation at SNPs between ancestral populations. Hence, we performed comparative study of allelic frequencies between the two ancestral populations and fine-mapping analysis to assess the potential presence of multiple causal variants. First, we computed allele frequency differences (ΔDAF)^50^ at all SNPs to measure the allelic heterogeneity between European and African population groups from the 1,000 Genomes Project (Phase III)^38^. The ΔDAF measures were previously found to be highly correlated with Weir and Cockerham’s F_ST_ in the 1,000 Genomes sample^51^. We assessed cases of extreme ΔDAF in the regions of interest by comparing the observed value against the threshold proposed by Colona et al.^24^ In brief, that study grouped 36.8 million variants in African and European populations from the 1,000 Genomes Project^38^ into bins of non-overlapping sets of 5,000 variants and derived the distribution of the maximum ΔDAF for further comparisons. Second, we applied a Bayesian method implemented in the software package FINEMAP^52^ to estimate the posterior probability of each single SNP interaction to be causal, conditionally on in-sample linkage disequilibrium pattern. FINEMAP implements a shotgun stochastic search algorithm to efficiently explore the most likely causal configurations, and we ran FINEMAP with the default parameters on the maximum number of causal SNPs (5), prior probabilities on the number of causal SNPs and the prior probabilities of a single SNP to be causal.

## Results

### Participants characteristics and description of traits

We used a dataset of 3,300 African American research volunteers from the Genetic Epidemiology of COPD (COPDGene) study^23^. Self-identified non-Hispanic African Americans and non-Hispanic European Americans between 45 and 80 years of age with a history at least 10 pack-years of smoking were enrolled from 21 medical centers across the US. Details on phenotyping, genome-wide genotyping of the cohort and inference of local ancestry are provided in Methods and **Supplementary Material**.

Individual characteristics of the COPDGene dataset are described in **Table 3** for the whole sample and stratified by current smoking status: non-smokers, moderate smokers (1-14 cigarettes per day) and heavy smokers (>14 cigarettes per day). The three smoker groups have similar overall characteristics, although there are some differences due to the COPDGene study enrollment protocol: the current smokers are younger and have higher proportions of males; non-smokers have higher numbers of accumulated pack-years. After stringent quality control procedure (**Supplementary Figures S8-S9** and **Supplementary Table S7**), we observed a total of 30,043 local ancestry segments. The distribution of proportions of local African ancestry (averaged across individuals) is roughly uniform along the genome (**Figure 1a**), confirming that the local ancestry data were free from artifacts. The proportion of global African ancestry ranged between 26.3% and 99.8% across individuals with an average of 80.3% (**Figure 1b**). We confirmed that individuals with higher proportions of African ancestry tend to have lower pulmonary function at a highly significant level (P<0.001, **Supplementary Table S4**) for all the traits except FEV_1_/FVC. We also assessed the variance heterogeneity among the smoking groups (not current smokers; moderate current smokers and heavy current smokers) by its explicit modeling as random effects (Equation 2). Both exploratory data analysis (**Supplementary Figure S7**) and formal statistical tests (P<0.0001 for all traits; **Supplementary Table S5**) highlighted differences among these three groups not only by mean, but also by variance (the group of non-smokers has the largest variance).

**Figure 1.**
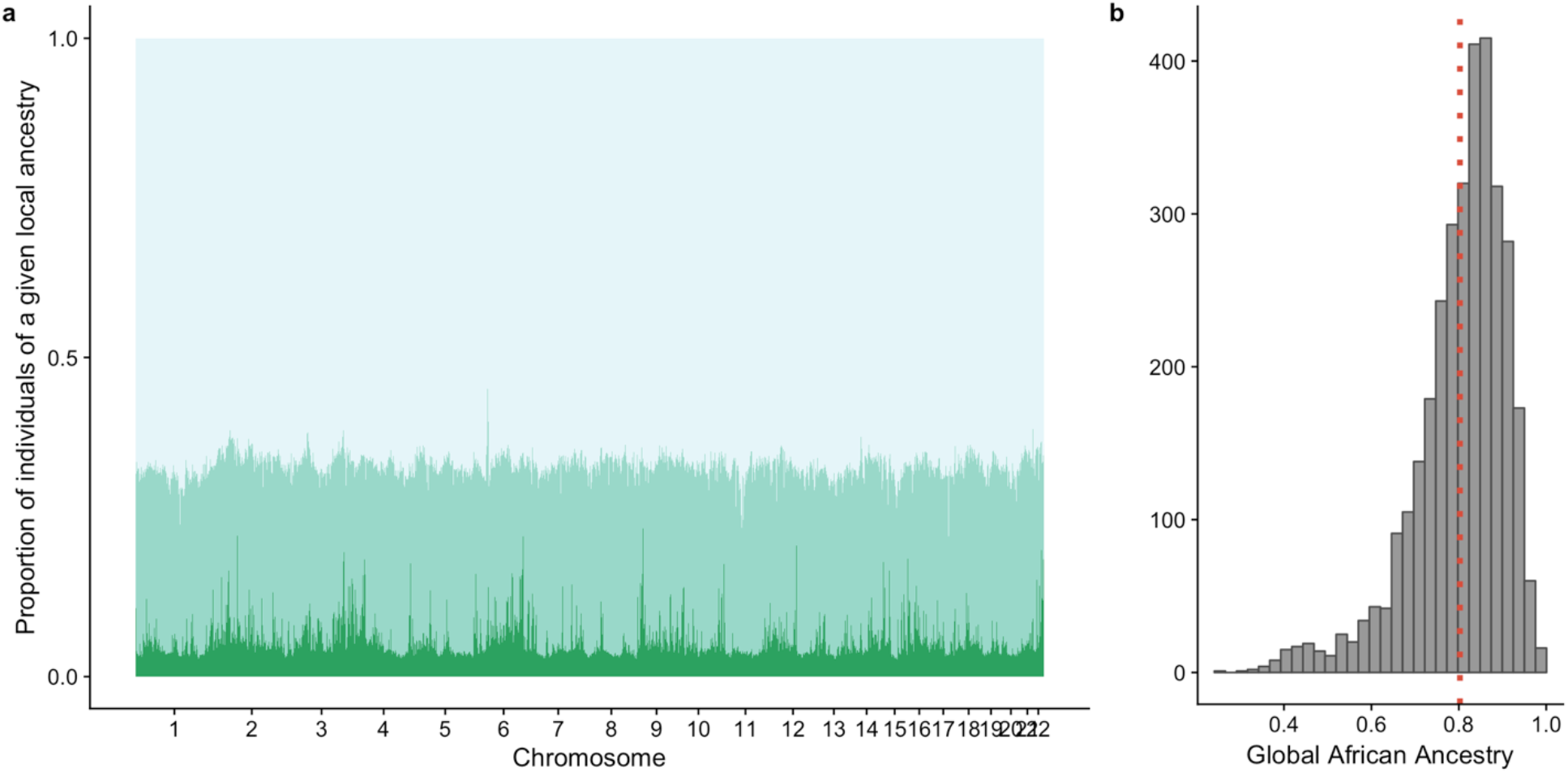
The African ancestry of African American participants of the COPDGene project. (a) The distribution of local ancestry is plotted by physical position in the genome on X axis. For each local ancestry segment the proportions of individuals with two African chromosomes (light green color), one African chromosome (green color) and no African chromosomes (dark green color) are presented on Y axis. (b) The distribution of the global African ancestry among 3,300 African-American individuals in the COPDGene study is shown. The vertical red dashed line depicts the mean value, 0.803.

**Table 1.** Characteristics of African American participants in the COPDGene project.

|  | Former smokers | Current moderate smokers | Current heavy smokers | All individuals |
| --- | --- | --- | --- | --- |
| Current cigarettes per day | 0 | 1-14 | >14 | – |
| Number of individuals (%) | 657 (20%) | 1,261 (38%) | 1,382 (42%) | 3,300 (100%) |
| Age enrolled | 60 (9) | 54 (6) | 53 (6) | 55 (7) |
| Gender, % male | 48% | 53% | 63% | 56% |
| Body-mass index | 30 (7) | 29 (7) | 29 (6) | 29 (7) |
| Pack-years smoked | 39 (22) | 32 (20) | 44 (21) | 38 (22) |
| Smoking duration, years | 34 (10) | 37 (8) | 36 (8) | 36 (8) |
| Global African ancestry, % | 79% (12%) | 81% (10%) | 80% (11%) | 80% (11%) |

*Dataset characteristics are presented for each smoking status separately and for all individuals. Values for a quantitative characteristic are given as the mean (standard deviation).*

**Table 2.** Top local ancestry segments-smoking interactions

| Locus | Ancestry segment | Exposure | Multi-trait P | Top single-trait P | Top trait |
| --- | --- | --- | --- | --- | --- |
| 11p15.2-3 | 12,075,829-12,845,835 | Current smoker | $2.8 \times 10^{-5}$ * | $5.8 \times 10^{-6}$ * | FEV<sub>1</sub> % predicted |
| 2q37.3 | 238,143,387-238,769,892 | Current heavy smoker | $2.9 \times 10^{-5}$ * | $2.5 \times 10^{-6}$ * | FEV<sub>1</sub> |
| 13q12.3-13.1 | 31,623,839-32,256,475 | Current heavy smoker | $3.4 \times 10^{-5}$ | 0.0052 | FVC |
| 11q21 | 94,360,812-94,825,729 | Current heavy smoker | $5.1 \times 10^{-5}$ | 0.0028 | FEV<sub>1</sub> |
| 7p15.2-3 | 25,133,849-26,371,279 | Current heavy smoker | $1.3 \times 10^{-4}$ | $2.82 \times 10^{-4}$ | FEV<sub>1</sub> % predicted |
| 8q21.13 | 81,871,222-82,335,354 | Current heavy smoker | $2.0 \times 10^{-4}$ | 0.24 | FVC |
| 1q44 | 248,020,448-249,208,153 | Current smoker | $3.2 \times 10^{-4}$ | 0.0029 | FEV<sub>1</sub>/FVC |

*Top signals from two admixture mappings of ancestry-smoking interactions, where environment exposure is either current smoker or current heavy smoker. Genome-wide significant association signals with p-value below the effective Bonferroni threshold $0.05/1,635 = 3.06 \times 10^{-5}$ are denoted with the “\*” mark, where 1,635 is the effective number of tests estimated by the eigenMT method<sup>18</sup>.*

**Table 3.** Top SNP-smoking interactions in regions identified by admixture mapping

| Locus | SNP | Position | Type | Gene | Anc. Z<sup>a</sup> | Anc. P<sup>a</sup> | SNP Z<sup>b</sup> | SNP P<sup>b</sup> | Allele | Freq<sub>CEU/YRI</sub> |
| --- | --- | --- | --- | --- | --- | --- | --- | --- | --- | --- |
| 11p15.2-3 | rs933920 | 12,481,110 | intronic | PARVA<sup>26,27</sup> | 4.32 | $1.6 \times 10^{-5}$ | 2.91 | 0.0036 | C/T | 0.99/0.89 |
| 11p15.2-3 | rs4553350 | 12,759,834 | intronic | TEAD1 | 4.20 | $2.7 \times 10^{-5}$ | -2.83 | 0.0046 | C/T | 0.55/0.95 |
| 2q37.3 | rs7569427 | 238,413,338 | intronic | MLPH | 4.63 | $3.7 \times 10^{-6}$ | -2.33 | 0.020 | A/G | 0.94/0.28 |
| 2q37.3 | rs2280289 | 238,483,729 | missense | RAB17<sup>28,29</sup> | 4.71 | $2.5 \times 10^{-6}$ | 2.10 | 0.036 | C/T | 0.84/0.20 |
| 13q12.3-13.1 | rs1535532 | 32,114,398 | intergenic |  | 2.18 | 0.030 | 2.32 | 0.020 | A/G | 0.67/0.62 |
| 11q21 | rs11020968 | 94,602,414 | missense | AMOTL1<sup>31</sup> | 2.77 | 0.0056 | -2.11 | 0.035 | A/T | 0.84/1.00 |
| 7p15.2-3 | rs10270076 | 25,363,037 | intergenic | | 3.31 | $9.3 \times 10^{-4}$ | -2.83 | 0.0046 | A/G | 0.57/0.29 |
| 8q21.13 | rs7000934 | 82,123,005 | intergenic |  | -1.17 | 0.24 | -2.38 | 0.017 | C/T | 0/0.15 |
| 1q44 | rs7533237 | 248,298,552 | intergenic |  | -2.54 | 0.011 | -2.83 | 0.0046 | G/T | 0/0.11 |

*Top SNP-smoking interaction signals for 7 loci identified by admixture mapping. Two top SNPs are listed for the two genome-wide significant loci (the first four rows). The reference allele frequencies in two CEU (European) and YRI (African) populations from the 1,000 Genomes Projects are presented for each SNP in the last two columns.*
<sup>a</sup>Z-scores and p-values are reported for the effects of local ancestry-exposure interactions.
<sup>b</sup>Z-scores and p-values are reported for the effects of SNP-exposure interactions.

### Mixed model to account for population structure and variance heterogeneity

To address the methodological question of performing a test of local ancestry-exposure interaction in admixed population, we chose a data-driven approach where the robustness of candidate models was assessed using the genomic control parameter (*λ*) and the overall shape of the standard quantile-quantile plot (QQ-plot) of the −log_10_(p-value). We started with the simplest model testing the marginal effect of local ancestry, and incrementally added terms necessary for robust testing of local ancestry-exposure interaction, our main parameter of interest.

We proposed to use a linear mixed model (LMM) including up to four random effects: *u_m_, u_i_, u_h_*, and *u_c_*, capturing structure due to shared local ancestry, local ancestry-smoking interaction, smoking status, and recruitment medical center, respectively. We added each of these components into the model through a step-wise procedure, assessing their relative contribution in various combinations. All variance-covariance matrices for the random effects observed in real data are illustrated in **Supplementary Figure S10**, while additional details of the model selection are provided in **Supplementary Material**.

We used the FEV_1_% predicted phenotype as an illustrative scenario and started with a marginal association model for single-trait admixture mapping (**Figure 2a**) (i.e. without ancestry-environment interaction; see also **Supplemental Notes** and **Supplementary Table S1**) including only *u_c_* as there was strong heterogeneity in the distribution of traits by medical center. Marginal local ancestry statistics in this initial model showed substantial inflation (*λ* = 1.19). We further accounted for correlation by genome-wide local ancestry across individuals, which was done by modeling the ancestral relationship matrix (ARM) (the *u_m_* random effect in Equation 2). The addition of the ARM component substantially mitigated the inflation (*λ* = 1.04, **Figure 2b**). We next considered adding the *u_h_* random effect that captured substantial portion of variance of modeled traits (**Supplementary Table S5**). This additional parameter did not impact the overall distribution of p-values (*λ* = 1.03), but resulted in an improved power of admixture mapping (the average of top statistics with P< 0.001 increased from 11.95 to 12.41), likely due to reduced amount of the residual variance after modeling the heterogeneity.

**Figure 2.**
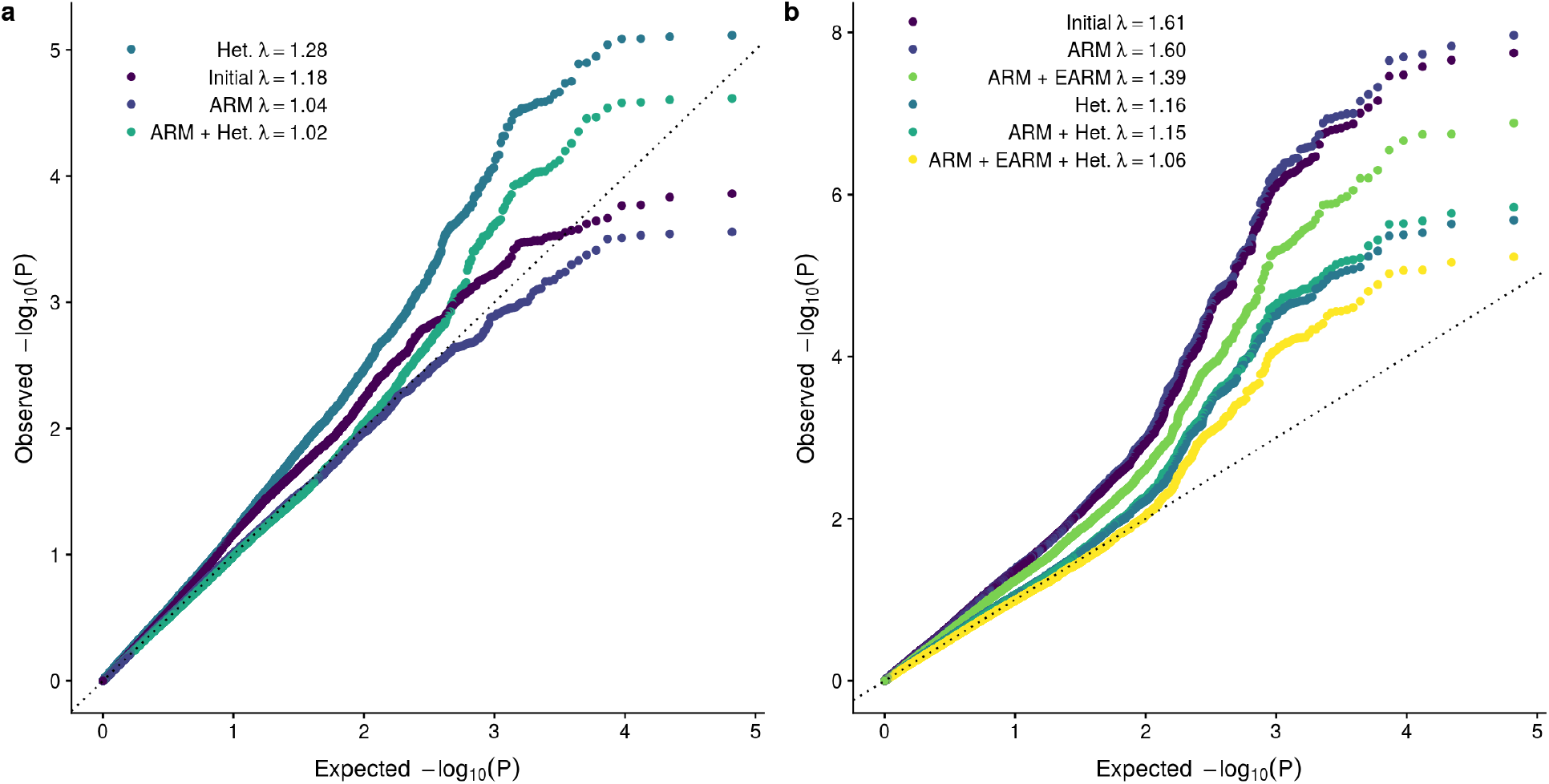
Robustness of mixed-model admixture mapping assessed by quantile-quantile (QQ) plots. The linear mixed model is examined through different combinations of random genetic and heterogeneity effects denoted in equations as u_m_, u_i_ and u_h_, while labeled on the plots as ARM, EARM and Het. (see Equation 1 and **Supplementary Table S1**). Admixture mapping is conducted for FEV_1_ % predicted phenotype, and current smoker status is used in evaluation of ancestry-smoking interactions. (a) When testing marginal effects of ancestry, the distribution of the test statistics is not inflated only when the genetic random effect (ARM) is presented in the model (*λ* = 1.04 or 1.02). (b) When testing ancestry-smoking interaction effects, all three random effects (ARM, EARM and Het.) are necessary to mitigate the inflation (*λ* = 1.06).

We then expanded the LMM to address the model components related to testing local ancestry-smoking interactions (Equation 1 and **Supplementary Table S1**). Following the exploration of the marginal model, we examined the role of four random effects by conducting admixture mapping for a single trait and the current smoker exposure in different model configurations (**Figure 2b**). Overall, the association test statistic was substantially inflated (*λ* = 1.61) if none of the three components were modeled. Including the components separately in the model, either the heterogeneity component, one genetic component with ARM or two genetic components, was not sufficient to fix the inflation (*λ* = 1, 16, 1.6 and 1.39, respectively). The association test statistic was well-behaved only when all three components were included (*λ* = 1.06). Hence, our final model (Equation 2) for admixture mapping of local ancestry-smoking (gene-environment) interaction included three additional random effects that capture structures due to shared local ancestry, local ancestry-smoking interaction and outcome heterogeneity across smoking groups.

### Local ancestry-smoking interactions detected by admixture mapping

We conducted admixture mapping of local ancestry-environment interaction for all five spirometric traits considering the two binary exposures independently (**Supplementary Figures S1 and S2**). Single-trait statistics showed limited inflation for all analyses (*λ* = 0.93-1.09), except for FEV_1_/FVC and current smoker exposure (*λ* = 1.12, **Supplementary Figures S4 and S5**). For each exposure, single traits results were combined to form a multi-trait test (**Figure 3**). The two multi-trait analyses show a well-controlled type I error rate (*λ* = 0.98, 1.02, **Supplementary Figures S6**). Following the *eigenMT* approach^18^, we estimated the number of effective tests to be 1,635 (**Supplementary Material**) and, thus, reduced the multiple-testing burden from the Bonferroni threshold (0.05/30,043 = 1.51×10^−6^ to the effective Bonferroni threshold (0.05/1,635 = 3.06×10^−5^). We also considered the suggestive Bonferroni threshold (0.5/1,635 = 3.06×10^−4^) to select additional candidate loci of interest.

**Figure 3.**
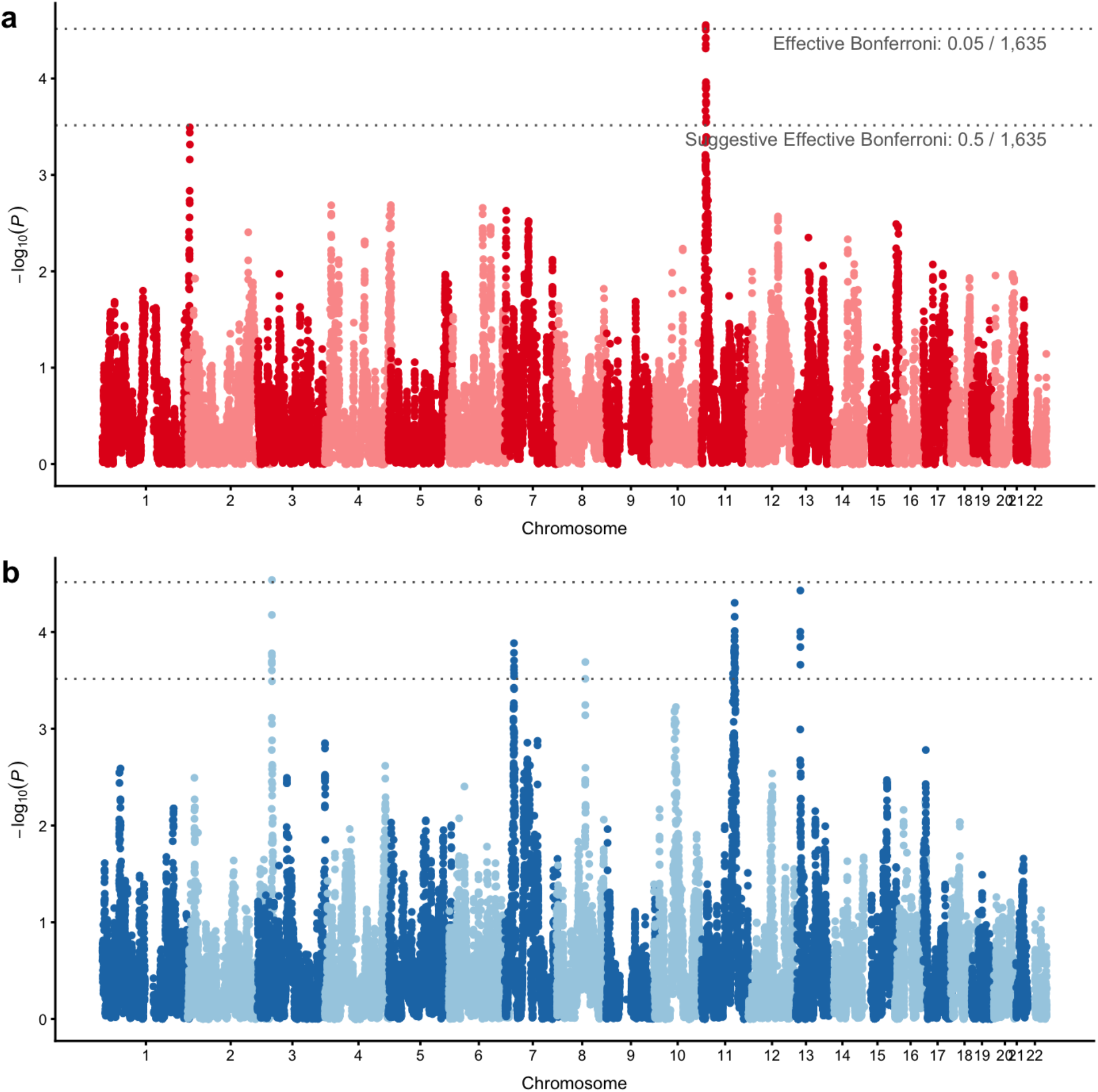
Admixture mapping identifies two genome-wide significant and five suggestive loci of local ancestry-smoking interactions. Manhattan plots show results of two admixture mappings of ancestry-smoking interactions, where smoking is one of two binary variables: (a) current smokers vs. non-smokers, and (b) current moderate smokers vs. current heavy smokers. The multivariate test joins the single-trait test statistics from five traits under the composite null hypothesis of no association and provides the multi-trait p-values. Horizontal lines depict the effective Bonferroni threshold (0.05/1,635 = *3.06×10^−5^*) and the effective suggestive Bonferroni threshold (0.5/1,635 = *3.06×10^−4^*), where 1,635 is the effective number of tests estimated by the eigenMT method^18^.

We identified two genome-wide significant and five suggestive interaction association signals (**Table 2**). The first genome-wide significant locus in the region chromosome 11p15.2-3 spanned 11 ancestry segments of average length 37Kb and 17 SNPs. The top ancestry segment Chr11: 12,341,061-12,373,680 of length 32,619b and 21 SNPs had a multi-trait positive interaction effect with current smoker exposure with multi-trait Z = 5.34 and P = 2.79×10^−5^. The signal was driven by all single-trait associations, where all p-values passed the suggestive Bonferroni threshold. The top single-trait association for FEV_1_ % predicted showed Z = 4.53 and P = 5.86×10^−6^. That top single-trait ancestry segment was 47kb away from the top multi-trait one, but the two segments were highly correlated after adjusting for the global ancestry (the Pearson correlation coefficient equaled 0.98). The second genome-wide significant locus in chromosome 2q37.3 included 11 ancestry segments of length 32Kb and 9 SNPs on average. The strongest multi-trait and top single-trait (FEV_1_) association signals were both localized in the same ancestry segment Chr2: 238,430,224-238,486,767 of length 56,543b and 21 SNPs. The positive interaction effects with the current heavy smoker exposure showed Z = 5.34 and *P* = 2.90×10^−6^, and Z = 4.53 and P = 2.50×10^−6^ for multi-trait and top single-trait associations, respectively.

The other five suggestive loci (**Table 2**) were mostly detected in the interaction admixture mapping with current heavy smoker exposure (13q12.3-13.1, 11q21, 7p15.2-3 and 8q21.13), except one locus 1q44 from the mapping with current smoking exposure. In contrast to the genome-wide significant loci, multi-trait association signals were much stronger than top single-trait signals for all suggestive loci (the difference in p-values was several orders of magnitude for some tests).

### Genotype-smoking interactions reveal differentiated genetic variants

For each region showing at least suggestive significance in the multi-trait admixture mapping analysis, we assessed potential interactions of single SNPs available in the region around the top admixture signal. We conducted association analysis for a total of 888 SNPs available across the 7 regions lying within ancestry segments (the average number of SNPs per region was 126.9, and the average number of SNPs per segment was 13.7). Here, we focused on the single trait showing the largest association signal in the admixture mapping. Overall, none of these SNPs passed a stringent Bonferroni correction threshold accounting for all SNPs tested in each region. However, the top SNPs especially in the genome-wide significant loci helped to localize the association signal (**Table 3**). In the first genome-wide significant region 11p15.2-3 (**Figure 4**), the top SNP rs933920 (P = 0.0036) is an intronic variant in the *PARVA* gene (MIM 608120). In the second genome-wide significant region 2q37.3 (**Figure 4**), the first top SNP rs7569427 (P = 0.02) was an intronic variant in the *MLPH* gene (MIM 606526), and second top SNP rs2280289 (P = 0.036) was a missense variant in the *RAB17* gene (MIM 602206).

**Figure 4.**
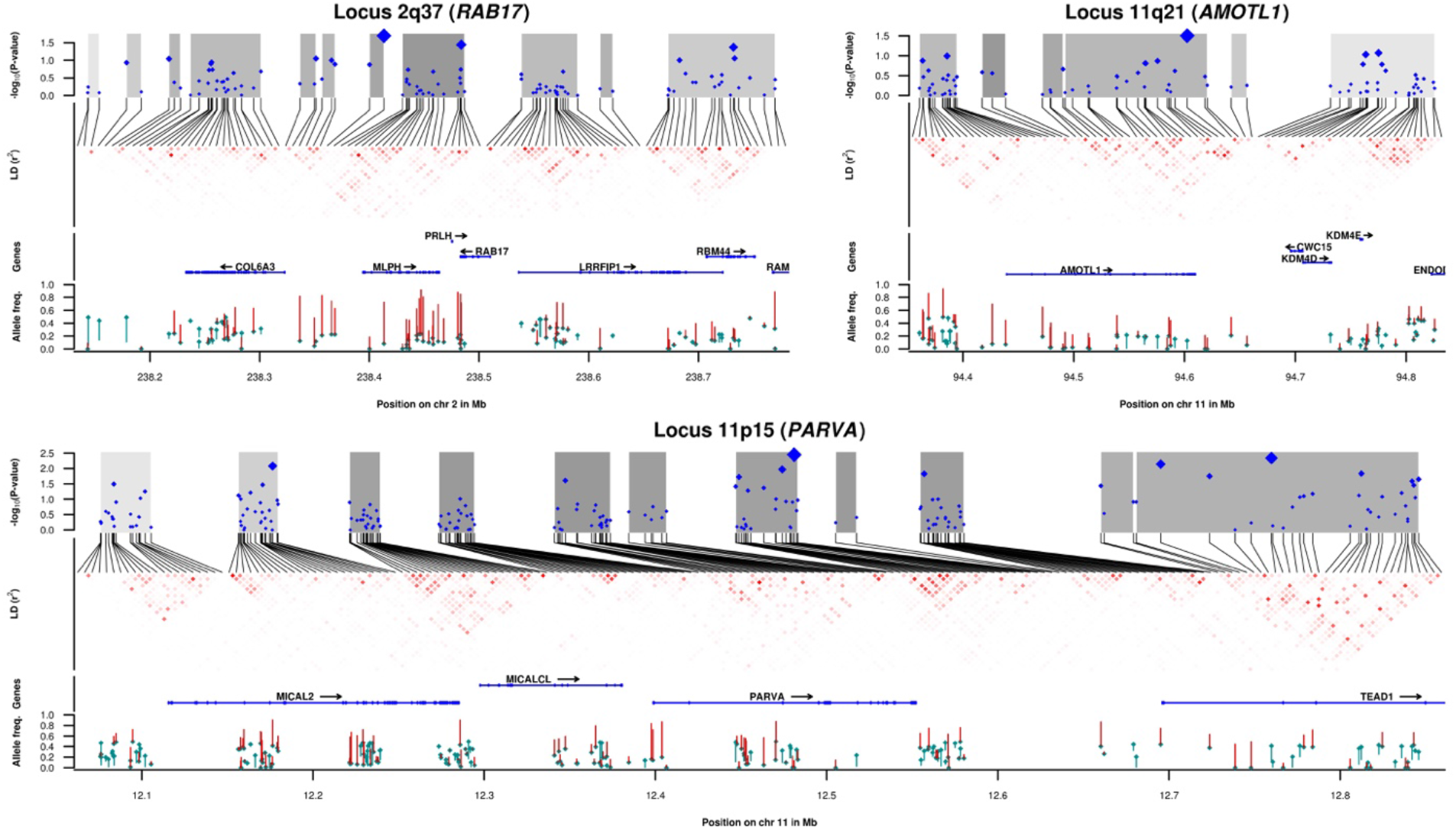
Detailed regional association plots for three selected loci 11p15.2-3, 2q37.3 and 11q21. From top to bottom, each panel shows (i) the regional association plot; (ii) linkage disequilibrium pattern; (iii) annotated protein-coding genes; (iv) trans-continental differences in allele frequency. (i) The Y axis represents −log_10_(P) of SNP association tests using a single phenotype most strongly associated with the ancestry segment in the locus. The shaded area represents the strength of local ancestry association in the multi-trait admixture mapping with stronger associations painted by darker shades of grey. The blue diamonds represent the Bayes factors for assessing the evidence that a SNP is causal estimated by FINEMAP; the bigger the diamond the higher the Bayes factor. (ii) The r^2^-based LD heatmap is built using genotypes of the COPDGene study, and the gradient of red is proportional to the r^2^. (iii) Protein-coding genes are queried from grch37.ensembl.org (iv) Allele frequencies are estimated in the 1,000 Genomes Project for European and African populations. For each SNP in an ancestry segment, the cyan indicates the frequency of the European minor allele variant, while a vertical segment connects the European and African frequencies of the allele. The segments are colored according to the direction of the difference: red when the African frequency is higher than the European frequency, or green for lower African frequency.

To assess the level of allelic heterogeneity of SNPs in these identified regions, we computed the allele frequency differences (defined as ΔDAF in Materials and Methods). The SNPs exhibited high levels of heterogeneity for all regions (ΔDAF in 0.62 and 0.78). Overall, 3 out of our 7 loci (1q44, 2q37.3 and 8q21.13) matched the 0.7 threshold proposed by Colonna *et al.*^24^, defining the 1% of the genome displaying the most extreme differentiation across populations. These differences in minimum allele frequency (MAF) between European and African ancestries can also be visually assessed on **Figure 4** and **Supplementary Figures S12-15**. Finally, we also evaluated the hypothesis of multiple causal SNPS per region using the FINEMAP software^21^, but we were not able to find any strong evidence for multiple causal SNPs (**Supplementary Table S6**).

### Replication of association signals in European GWASs

We performed replication analyses of association signals detected at individual SNP-level and gene-level, for the seven loci reported in **Tables 2–3**. We considered two large studies of pulmonary phenotypes and COPD conducted in individuals of European ancestry: the CHARGE consortium (N = 50,047) which had genome-wide summary results for SNP-by-smoking interaction^53^; and the most recent and largest meta-analysis of UK Biobank and SpiroMeta cohorts (COPD cases = 35,735, COPD controls = 222,076; N = 400,102 for pulmonary function phenotypes) which provides an up-to-date list of variants with genome-wide significant marginal genetic effect.^54,55^.

Matching our nine top SNPs with the CHARGE consortium results^53^, we found that three were missing, being rare or monomorphic in European population. Interaction effects of three of out the six remaining SNPs were replicated at the nominal significance level (P <0.05) (**Supplementary Table S8**) in SNP-smoking interaction screening of FEV_1_/FVC, where packs-years was used as a proxy for heavy smoking. We also assessed the joint SNP and SNP-smoking association reported in the CHARGE consortium^53^ in all seven loci. Although, no loci display genome-wide significant association, we observed that all but one showed at least one SNP with association signal at the level of P< 0.0001 (the strongest signal for rs10202058 with P = 1.92 x10^−6^ in the 2q37.3 locus).

We next assessed the presence of marginal genetic effect at our seven loci using the aforementioned meta-analysis of pulmonary phenotypes^54^ and COPD^55^. Although our marginal signals of top local ancestry segments and SNPs were weak (**Supplementary Table S10-11**), we observed that nine genome-wide significant SNPs from GWASs^54,55^ were located less than 1Mb away from the three loci 2q37.3, 11p15.2-3 and 7p15.2-3 (**Supplementary Table S9**). In particular, two SNPs rs80145403 and rs80145403 were within the same *PARVA* and *TEAD1* genes in Chromosome 11 as in our SNP-smoking interaction analysis (**Table 3**). Notably, five SNPs come from two loci that each has three distinct signals estimated by the conditional analysis^55^. Such a scenario with multiple SNPs driving either marginal or interaction association is beneficial for our ancestry-based approach to detect gene-environment interactions, and, thus, may explain the signal overlap at the gene-level for two 11p15.2-3 and 2q37.3 loci.

## Discussion

Broadening the diversity of ethnicities in genetic analysis can provide important information for disease pathogenesis. Leveraging local ancestry through admixture mapping could improve power to discover marginal genetic and gene-environment effects, although the technical and statistical challenges still remain. To address these challenges, we introduced a multi-component linear mixed model and empirically demonstrated its robustness in admixture mapping on real data in 3,300 African American participants in the COPDGene study. We detected two genome-wide significant and five suggestive loci showing smoking-dependent effects of local ancestry on spirometric lung function phenotypes.

While the functional effects of variants in the identified genomic regions is unknown, these regions harbor genes previously known for traits related to smoking. The top SNP rs93392 in the first genome-wide significant locus 11p15.2-3 (P = 2.79×10^−5^) is located within the *PARVA* gene, which produces a focal adhesion protein^25^. Two previous studies reported that this gene was differentially methylated in small airway epithelium^26^ and buccal mucosa^27^ when stratified by current smoking status. The second top SNP rs2280289 for the second genome-wide significant region on 2q37.3 (P = 2.90×10^−5^) is a missense mutation in *RAB17,* which was previously associated with a smoking cessation genotype success score^28,29^. As for the most relevant result found among the five suggestive loci, the top SNP rs11020968 in the locus 11q21 (P = 5.0×10^−5^) is a missense mutation for angiomotin-like protein 1 *(AMOTL1;* MIM 614657), a tight junction protein hypothesized to play a role in COPD through endothelial tight junction permeability, and whose expression is affected by cigarette smoking^30^. Additional genes around other loci may warrant further investigation.

We further attempted to evaluate whether SNPs within these identified regions show multiple-SNP effects and exhibit high allelic differentiation, as our previous work on gene-gene interaction admixture mapping suggested this kind of genetic architecture^18^. Overall, allele frequency heterogeneity between European and African ancestries was very strong and persistent in the identified regions. Although our fine-mapping analysis did not show evidence for multiple causal variants, SNP-smoking interaction analysis is known to have limited power^31^; thus, we cannot rule out the possibility of multiple causal variants. Indeed, the conditioning on the primary signal of local ancestry is likely to decrease the statistical power to detect interactions at the SNP level even in larger samples^31^. Alternative methods, such as jointly modeling ancestry and genotype association signals, might help to overcome this limitation^32^.

Our methodological contributions to admixture mapping are multiple. First, we extended the concept of genetic relationship matrix originally proposed to control population structure in GWAS^33^: the ancestry relationship matrix (ARM) was similarly computed on local ancestry data and further used in association tests. Second, we adopted the population stratification approach recently designed specifically for GWAS of gene-environment interactions^34^: two matrices, the standard ARM, but also a second environmental ARM or EARM, were essential to control for spurious association results when testing local ancestry-environment interactions. Finally, we modeled the outcome heterogeneity among groups stratified by environmental exposure. Modeling this heterogeneity increased the power because of reduced residual phenotypic variance (up to 65%); and substantially decreased the inflation of interaction test statistic.

Our study also has limitations. COPDGene is one of the largest studies of African-American smokers, with a high proportion of subjects with COPD, which makes suitable replication cohorts challenging. Nevertheless, we were able to reproduce some of the top SNP-smoking interactions in the CHARGE consortium^53^ at nominal significant level. More importantly, our study identified 3 loci 2q37.3, 11p15.2-3 and 7p15.2-3 using a dataset of only 3,300 African-American individuals, while the same loci only passed the genome-wide significance threshold of standard univariate association in an independent replication cohort including up to 400,000 individuals^54,55^. Further, the method for admixture mapping can also be optimized. When conducting association analysis, excluding the local ancestry segment under testing from the ARM construction will be able improve power^35^, but is computationally more burdensome. We attempted a more efficient out-of-chromosome strategy commonly applied in GWAS^36^, but we observed fairly inflated test statistics (data not shown).

In conclusion, our study reports a powerful approach for gene-environment interaction association studies, leveraging the unique genetic architecture of complex traits measured in recently admixed populations. The proposed statistical model has shown to be robust to population structure and outcome variance heterogeneity. In our application to the COPDGene study, we have found two genome-wide significant local ancestry-smoking interactions of lung function phenotypes that would have been missed in standard single SNP interaction analyses. Overall, our findings provide additional evidence of the importance of ethnic diversity in genetic clinical studies.

## Acknowledgements

This project was supported by NHGRI R21HG007687 (A.Z., H.A. and M.H.C.), and NHLBI R01HL113264 and R01HL137927 (M.H.C.) The COPDGene study (NCT00608764) is supported by NHLBI R01 HL089897 and R01 HL089856, and the COPD Foundation through contributions made to an Industry Advisory Board composed of AstraZeneca, Boehringer Ingelheim, Novartis, Pfizer, GlaxoSmithKline, Siemens, and Sunovion. M.H.C. has received grant support from GSK. The content is solely the responsibility of the authors and does not necessarily represent the official views of the National Heart, Lung, and Blood Institute or the National Institutes of Health. Additional information can be found in the Supplemental Material.

## Code and Data Availability

Code used here for admixture mapping, https://gist.github.com/variani/a28c18797c39a62bacab587e6e708529 The 1,000 Genomes Project (Phase III, version 5), http://csg.sph.umich.edu/abecasis/mach/download/1000G.Phase3.v5.html LAMP-LD software for ancestry inference, http://bogdan.bioinformatics.ucla.edu/software/lamp/ Gaston R package for mixed models, https://cran.r-project.org/package=gaston Public GWAS summary-statistics from the CHARGE consortium, the dbGaP database, study accession phs000930.v3.p1, https://www.ncbi.nlm.nih.gov/projects/gap/cgi-bin/study.cgi?study_id=phs000930.v3.p1 OMIM, http://www.omim.org

