## Supplementary Material for "Mixed-model admixture mapping identifies smoking-dependent loci of lung function in African Americans"

**COPDGene® Investigators – Core Units**

*Administrative Center*: James D. Crapo, MD (PI); Edwin K. Silverman, MD, PhD (PI); Barry J. Make, MD; Elizabeth A. Regan, MD, PhD

*Genetic Analysis Center*: Terri Beaty, PhD; Ferdouse Begum, PhD; Peter J. Castaldi, MD, MSc; Michael Cho, MD; Dawn L. DeMeo, MD, MPH; Adel R. Boueiz, MD; Marilyn G. Foreman, MD, MS; Eitan Halper-Stromberg; Lystra P. Hayden, MD, MMSc; Craig P. Hersh, MD, MPH; Jacqueline Hetmanski, MS, MPH; Brian D. Hobbs, MD; John E. Hokanson, MPH, PhD; Nan Laird, PhD; Christoph Lange, PhD; Sharon M. Lutz, PhD; Merry-Lynn McDonald, PhD; Margaret M. Parker, PhD; Dandi Qiao, PhD; Elizabeth A. Regan, MD, PhD; Edwin K. Silverman, MD, PhD; Emily S. Wan, MD; Sungho Won, Ph.D.; Phuwanat Sakornsakolpat, M.D.; Dmitry Prokopenko, Ph.D.

*Imaging Center*: Mustafa Al Qaisi, MD; Harvey O. Coxson, PhD; Teresa Gray; MeiLan K. Han, MD, MS; Eric A. Hoffman, PhD; Stephen Humphries, PhD; Francine L. Jacobson, MD, MPH; Philip F. Judy, PhD; Ella A. Kazerooni, MD; Alex Kluiber; David A. Lynch, MB; John D. Newell, Jr., MD; Elizabeth A. Regan, MD, PhD; James C. Ross, PhD; Raul San Jose Estepar, PhD; Joyce Schroeder, MD; Jered Sieren; Douglas Stinson; Berend C. Stoel, PhD; Juerg Tschirren, PhD; Edwin Van Beek, MD, PhD; Bram van Ginneken, PhD; Eva van Rikxoort, PhD; George Washko, MD; Carla G. Wilson, MS;

*PFT QA Center, Salt Lake City, UT*: Robert Jensen, PhD

*Data Coordinating Center and Biostatistics*, *National Jewish Health, Denver, CO*: Douglas Everett, PhD; Jim Crooks, PhD; Camille Moore, PhD; Matt Strand, PhD; Carla G. Wilson, MS

*Epidemiology Core*, *University of Colorado Anschutz Medical Campus, Aurora, CO*: John E. Hokanson, MPH, PhD; John Hughes, PhD; Gregory Kinney, MPH, PhD; Sharon M. Lutz, PhD; Katherine Pratte, MSPH; Kendra A. Young, PhD

*Mortality Adjudication Core:* Surya Bhatt, MD; Jessica Bon, MD; MeiLan K. Han, MD, MS; Barry Make, MD; Carlos Martinez, MD, MS; Susan Murray, ScD; Elizabeth Regan, MD; Xavier Soler, MD; Carla G. Wilson, MS

*Biomarker Core*: Russell P. Bowler, MD, PhD; Katerina Kechris, PhD; Farnoush Banaei-Kashani, Ph.D

**COPDGene® Investigators – Clinical Centers**

*Ann Arbor VA:* Jeffrey L. Curtis, MD; Carlos H. Martinez, MD, MPH; Perry G. Pernicano, MD

*Baylor College of Medicine, Houston, TX*: Nicola Hanania, MD, MS; Philip Alapat, MD; Mustafa Atik, MD; Venkata Bandi, MD; Aladin Boriek, PhD; Kalpatha Guntupalli, MD; Elizabeth Guy, MD; Arun Nachiappan, MD; Amit Parulekar, MD;

*Brigham and Women's Hospital, Boston, MA*: Dawn L. DeMeo, MD, MPH; Craig Hersh, MD, MPH; Francine L. Jacobson, MD, MPH; George Washko, MD

*Columbia University, New York, NY*: R. Graham Barr, MD, DrPH; John Austin, MD; Belinda D'Souza, MD; Gregory D.N. Pearson, MD; Anna Rozenshtein, MD, MPH, FACR; Byron Thomashow, MD

*Duke University Medical Center, Durham, NC*: Neil MacIntyre, Jr., MD; H. Page McAdams, MD; Lacey Washington, MD

*HealthPartners Research Institute, Minneapolis, MN*: Charlene McEvoy, MD, MPH; Joseph Tashjian, MD

*Johns Hopkins University, Baltimore, MD*: Robert Wise, MD; Robert Brown, MD; Nadia N. Hansel, MD, MPH; Karen Horton, MD; Allison Lambert, MD, MHS; Nirupama Putcha, MD, MHS

*Los Angeles Biomedical Research Institute at Harbor UCLA Medical Center, Torrance, CA*: Richard Casaburi, PhD, MD; Alessandra Adami, PhD; Matthew Budoff, MD; Hans Fischer, MD; Janos Porszasz, MD, PhD; Harry Rossiter, PhD; William Stringer, MD

*Michael E. DeBakey VAMC, Houston*, *TX*: Amir Sharafkhaneh, MD, PhD; Charlie Lan, DO

*Minneapolis VA:* Christine Wendt, MD; Brian Bell, MD

*Morehouse School of Medicine, Atlanta, GA*: Marilyn G. Foreman, MD, MS; Eugene Berkowitz, MD, PhD; Gloria Westney, MD, MS

*National Jewish Health, Denver, CO*: Russell Bowler, MD, PhD; David A. Lynch, MB

*Reliant Medical Group, Worcester, MA*: Richard Rosiello, MD; David Pace, MD

*Temple University, Philadelphia, PA:* Gerard Criner, MD; David Ciccolella, MD; Francis Cordova, MD; Chandra Dass, MD; Gilbert D'Alonzo, DO; Parag Desai, MD; Michael Jacobs, PharmD; Steven Kelsen, MD, PhD; Victor Kim, MD; A. James Mamary, MD; Nathaniel Marchetti, DO; Aditi Satti, MD; Kartik Shenoy, MD; Robert M. Steiner, MD; Alex Swift, MD; Irene Swift, MD; Maria Elena Vega-Sanchez, MD

*University of Alabama, Birmingham, AL:* Mark Dransfield, MD; William Bailey, MD; Surya Bhatt, MD; Anand Iyer, MD; Hrudaya Nath, MD; J. Michael Wells, MD

*University of California, San Diego, CA*: Joe Ramsdell, MD; Paul Friedman, MD; Xavier Soler, MD, PhD; Andrew Yen, MD

*University of Iowa, Iowa City, IA*: Alejandro P. Comellas, MD; Karin F. Hoth, PhD; John Newell, Jr., MD; Brad Thompson, MD

*University of Michigan, Ann Arbor, MI*: MeiLan K. Han, MD, MS; Ella Kazerooni, MD; Carlos H. Martinez, MD, MPH

*University of Minnesota, Minneapolis, MN*: Joanne Billings, MD; Abbie Begnaud, MD; Tadashi Allen, MD

*University of Pittsburgh, Pittsburgh, PA*: Frank Sciurba, MD; Jessica Bon, MD; Divay Chandra, MD, MSc; Carl Fuhrman, MD; Joel Weissfeld, MD, MPH

*University of Texas Health Science Center at San Antonio, San Antonio, TX*: Antonio Anzueto, MD; Sandra Adams, MD; Diego Maselli-Caceres, MD; Mario E. Ruiz, MD

### Supplementary notes

#### The COPDGene study

##### Phenotypes

Five spirometric measures of lung function were considered as quantitative phenotypes: forced expiratory volume in one second ($FEV_1$); forced expiratory volume in one second as a percent of predicted ($FEV_1$ % predicted); forced vital capacity (FVC); forced vital capacity as a percent of predicted (FVC predicted); and the ratio of forced expiratory volume in one second to forced vital capacity ($FEV_1$/FVC). $FEV_1$ is measured as the amount of air forcefully exhaled in one second. FVC is the total amount of air in a complete forced exhalation. Two phenotypes $FEV_1$ % predicted and FVC % predicted are calculated using race and gender-specific regression models based on age, age squared and height squared[1]. The ratio $FEV_1$/FVC is used to determine the presence of airflow obstruction (values below 0.7 are used for GOLD classification of COPD)[2]. Post-bronchodilator measurements obtained by the NDD EasyOne Spirometer (Zurich, Switzerland) were used in data analysis, where 3,260 study participants passed spirometric quality control.

##### Genotypes

The COPDGene individuals were genotyped using the Illumina Omni Express BeadChip with 733,200 single nucleotide polymorphisms (SNPs). The COPDGene genotyping quality control protocol for the whole sample of non-Hispanic African Americans and non-Hispanic European Americans is described elsewhere[3,4]. We downloaded the genotype data from the official COPDGene release (as of October, 2017), that contained 3,300 African American individuals and 684,255 SNPs that passed the quality control for GWAS (minimum allele frequency (MAF) >1%). Principal component analysis revealed none of the African American study participants to be excluded based on falling more than six standard deviations from the mean of the first two principal components. We also excluded SNPs that were unmatched or not presented or in the 1,000 Genomes reference panel[5]. As a result, 3,300 African American individuals 684,187 autosomal SNPs were available for further genetic admixture inference and association analysis.

##### Local ancestry inference with LAMP-LD

We first processed the output of the ancestry inference procedure performed by the LAMP-LD program[6]. We obtained estimates of having 0, 1 or 2 African chromosomes at each SNP position in the form of data matrix of 3,300 rows and 675,316 columns. When comparing to output from our previous ancestry analysis of the same dataset[3] (**Supplementary Table S7**), we observed a higher quality of the ancestry data (**Supplementary Figure S8**) because of the larger number of input SNPs (675,316 *vs.* 608,917) and the larger number of individuals in reference ancestral populations (207 in 1,000 Genomes *vs.* 120 in HapMap II). The later factor is established to have the major impact on the inference accuracy of the LAMP-LD algorithm[6].

The global African ancestry estimates in our analysis and previous work[3] were highly concordant (the Pearson correlation coefficient, r = 0.999, **Supplementary Figure S9**).

#### Statistical model for admixture mapping

Consider a quantitative phenotype stored in a $n$-dimensional vector $y$ and a $n \times m$ data matrix $Z$ of local ancestry segments, where $n$ is the number of individuals and $m$ is the number of local ancestry segments. Let $z_l$ be a column of matrix $Z$, a single local ancestry segment. Admixture mapping consists in testing each local ancestry segment ($z_l$) for correlation with phenotype ($y$) using linear regression model. When testing for gene-by-environment interaction, three variables are given in the model: the vector $z_l$, a vector $x_e$ of an environmental exposure, and an interaction term $z_g \times x_e$ computed as element-wise product of two vectors $z_l$ and $x_e$. Thus, admixture mapping of gene-by-environment interactions examines correlation of an interaction term ($z_g \times x_e$) with phenotype ($y$) in the presence of the other two variables ($z_l$ and $x_e$). Here, we first introduce a statistical model for admixture mapping (testing the effect of local ancestry on phenotype) and then describe its extension to admixture mapping of interactions (testing the effect of the interaction term on phenotype). Before presenting the two models, we introduce methods in genetic association studies to control for confounding due to population structure.

#### Confounding due to population structure

Admixture mapping performed on data collected from individuals of mixed ancestries inherently reveals systematic differences in allele frequencies between subgroups of the data sample, a phenomenon known as population structure. Population structure can be a confounder in genetic association studies, including admixture mapping, leading to both false positive and false negative associations. Admixture and population stratification are two models of population structure used in association studies.

Admixture is a result of mating among two or more isolated ancestry populations. Admixed individuals are characterized by ancestry proportions measured as genome-wide averages, referred to as the global ancestries. For example, African Americans represent two-way admixture and, thus, a single measure of the global African ancestry is enough to express admixed proportions for each individual. Including the global ancestries as covariates into linear model is the standard approach in admixture mapping, that allows to estimate the effect of local ancestry on phenotype and avoid confounding due to the global ancestries. Although the global ancestries capture the major signal due to admixture, the remaining phenotypic variance due to local ancestry needs to be modeled using the concept of genetic relationship matrices[7], which is explained below.

Population stratification is typically quantified and further incorporated into linear model by two widely used methods, principal component analysis and linear mixed models. Both methods make use of the realized genetic relationship matrix (GRM) computed on the genetic data matrix of SNPs (centered and scaled). The ancestral relationship matrix (ARM) is a similar construct for use in genetic data from admixed populations. This matrix can be estimated from the local ancestry data the same way and further incorporated into the linear mixed model to estimate the proportion of phenotypic variance attributable to additive genetic variance[7,8].

#### Testing the effect of local ancestry on phenotype

Consider the following single-trait liner mixed model that describes one of the five quantitative phenotypes $y$ in the dataset of 3,300 African American individuals from the COPDGene study:

$$y = X\beta + Zu_c + u_m + e (S1)$$

where $n = 3{,}300$ is the number of individuals, $p$ is the number of fixed effects that depends on a phenotype (**Supplementary Table S2**), $c = 21$ is the number of medical centers, $X_{n \times p}$ is an incidence matrix of fixed effects, $Z_{n \times c}$ is an incidence matrix of random effects of medical centers; $\beta$ is a vector of fixed effects, $u_c$ is a vector of random effects of medical centers, $u_m$ is a vector of genetic random effects of ancestry, and $e$ is a vector of the residuals errors. The random vectors $u_c$, $u_m$ and $e$ are mutually independent and multivariate normal distributed as $N(0, \sigma_c^2 I_{c \times c})$, $N(0, \sigma_m^2 A_{n \times n})$ and $N(0, \sigma_e^2 I_{n \times n})$, respectively. The matrices $I$ are the identity matrices and $A$ is the ancestral relationship matrix (ARM)[7,8]. Hence, the variance components $\sigma_c^2$, $\sigma_m^2$ and $\sigma_e^2$, as well as the effect sizes $\beta$, are model parameters to be estimated.

The matrix $A$ is computed by cross-product operation on 30,043 long ancestry segments, which were previously centered and scaled[8]. Let $z_{ij}$ represent the local ancestry for the $j$ individual at the $i$ of $m$ loci. This variable can take values 0, 1 or 2, which correspond to the number of chromosomes of African ancestry. Given $p = \frac{1}{2n}\sum_{j=1}^{n} z_{ij}$, the pairwise ancestral similarity between the $j$ and $k$ individuals is expressed as following:

$$A_{jk} = \frac{1}{m}\sum_{i=1}^{m}\frac{(z_{ji}-2p_i)(z_{ki}-2p_i)}{2p_i(1-p_i)} (S2)$$

Covariates stored in columns of the $X$ matrix include the intercept ($x_0$), phenotype-specific covariates, the global ancestry ($z_g$) and a local ancestry segment ($z_l$), which is to be tested for association with the phenotype $y$.

$$X = [x_0; \dots; z_g; z_l] (S3)$$

The test of association is the $\chi^2$-test with one degree of freedom, which statistic is computed using the estimated effect size for local ancestry $\hat{\beta}_{z_l}$ and its standard error $\hat{\sigma}_{z_l}$: $T = \hat{\beta}_{z_l}^2/\hat{\sigma}_{z_l}^2$.

**Testing the effect of interaction between local ancestry and smoking exposures**

The Equation S1 is expanded by adding both fixed and random effects terms (see also Equation 2 in the main text and Figure S10):

$$y = X\beta + u_m + u_i + u_h + u_c + e (S3)$$

$$X = [x_0; \dots; x_e; z_g; z_l; z_g x_e; z_l x_e] (S4)$$

The first additional term of random effects$u_i$is devoted to control for population structure in the settings when association test is performed for gene-environment (here, local ancestry-smoking) interactinos[14.] It has a variance-covariance matrix derived from ARM ($A_{nxn}$) and based on stratification by binary environmental exposure$x_e$. We refer to this matrix as EARM and further explain its construction. Define a matrix $D_{nxn}$such that its element $D_{ij}$is equal to 1 only if i^th^ and j^th^ entries in exposure vector $x_e$are in the same, i.e. both individuals i^th^ and j^th^ belong to the same exposed or unexposed strata. Otherwise, entries in $D_{nxn}$are zeros. Then the EARM matrix is calculated as element-wise multiplication of two matrices $D_{nxn}$and $A_{nxn}$[14].

The second additional term of random effects $u_h$is the standard way to model variance heterogeneity in outcome $y$across different individual groups (here environmental exposure stratas defined by $x_e$). The variance-covariance matrix of this term is a diagonal matrix $H_{nxn}$, where each group of individuals stratified by $x_e$ has its own variance component $\sigma_h^2$.

The test of association of interaction is the $\chi^2$-test with one degree of freedom, which statistic is computed using the estimated effect size for local ancestry -smoking interaction and its standard error, similarly as for the marginal effect of local ancestry in Equations S1 and S2.

#### Correction for multiple testing

To determine significant associations in admixture mapping, we computed the effective number of tests using the eigenMT methodology[9]. First, we observed that local ancestry segments exhibited strong correlation pattern (**Supplementary Figure S11**) in comparison to SNP data.

Next, we decreased the burden of multiple testing in admixed mapping by calculation of the effective number of ancestry segments. We followed the eigenMT method proposed for local expression quantitative trait locus (cis-eQTL) studies[9], which is more straightforward to apply and implement in our data analysis than other methods[10]. We estimated the sample correlation matrix on data matrix of ancestry segments by the Ledoit-Wolf regularized estimator[11] and calculated its eigen-value decomposition. The effective number of ancestry segments was computed as the minimum number of eigen-vectors that captures a given threshold for the proportion of variance explained. We set that threshold to 0.95. When estimating the effective number of ancestry segments, we projected out the global ancestry from the ancestry segment data. That operation follows from our association models (1) and (2) in the main text, where we test the effect of local ancestry covariate in the presence of the global ancestry included as another covariate. In other words, the tested effect is exclusively due to that variance in local ancestry that is complementary to the global ancestry under the linear model. That projection operation also allows us to separately estimate the number of effective local ancestry segments for each chromosome, if necessary, assuming that the intra-chromosomal correlation among local ancestry is mainly due to the global ancestry, which was projected out from local ancestry data.

#### Supplementary tables

**Supplementary Table S1. Linear mixed models used in step-wise model selection procedure.**

| Model Name | Model for marginal test ($\beta_l$) | Model for interaction test ($\delta_l$ |
|---|---|---|
| Initial | $y = C\beta_C + \beta_e x_e + \beta_g z_g + \beta_l z_l + u_c + e$ | $y = C\beta_C + \beta_e x_e + [\beta_g z_g + \delta_g z_g x_e] + [\beta_l z_l + \delta_l z_l x_e] + u_c + e$ |
| Initial + ARM | $y = C\beta_C + \beta_e x_e + \beta_g z_g + \beta_l z_l + u_c + u_m + e$ | $y = C\beta_C + \beta_e x_e + [\beta_g z_g + \delta_g z_g x_e] + [\beta_l z_l + \delta_l z_l x_e] + u_c + u_m + e$ |
| Initial + Het | $y = C\beta_C + \beta_e x_e + \beta_g z_g + \beta_l z_l + u_c + u_h + e$ | $y = C\beta_C + \beta_e x_e + [\beta_g z_g + \delta_g z_g x_e] + [\beta_l z_l + \delta_l z_l x_e] + u_c + u_h + e$ |
| Initial + ARM + Het | $y = C\beta_C + \beta_e x_e + \beta_g z_g + \beta_l z_l + u_c + u_m + u_h + e$ | $y = C\beta_C + \beta_e x_e + [\beta_g z_g + \delta_g z_g x_e] + [\beta_l z_l + \delta_l z_l x_e] + u_c + u_m + u_h + e$ |
| Initial + ARM + EARM | | $y = C\beta_C + \beta_e x_e + [\beta_g z_g + \delta_g z_g x_e] + [\beta_l z_l + \delta_l z_l x_e] + u_c + u_m + u_i + e$ |
| Initial + ARM + EARM + Het | | $y = C\beta_C + \beta_e x_e + [\beta_g z_g + \delta_g z_g x_e] + [\beta_l z_l + \delta_l z_l x_e] + u_c + u_m + u_i + u_h + e$ |

*Linear mixed models with different compositions of random effect are examined to provide a robust test of either marginal or interaction effect of local ancestry on quantitative trait. See the main text of manuscript describing the model notation.*

**Supplementary Table S2. Phenotype-specific covariates or fixed effects in linear mixed model.**

| Phenotype | Covariates |
|---|---|
| $FEV_1$ % predicted | age, gender, smoking (*) duration, cigarettes per day on average (log-transformed), current smoking status, current heavy smoking status (**), current smoking cigar status |
| $FEV_1$ | age, age squared, gender, height (cm), smoking duration, cigarettes per day on average (log-transformed), current smoking status, current heavy smoking status, current smoking cigar status |
| FVC % predicted | age, gender, smoking duration, cigarettes per day on average (log-transformed), cur- rent smoking status, current heavy smoking status, current smoking cigar status |
| FVC | age, age squared, gender, height (cm), smoking duration, cigarettes per day on aver- age (log-transformed), current smoking status, current heavy smoking status, current smoking cigar status |
| $FEV_1$ /FVC | age, age squared, gender, smoking duration, cigarettes per day on average (log-transformed), current smoking status, current heavy smoking status |

*(*) We refer to smoking cigarettes when saying smoking status. (**) The current heavy smoking status is defined as smoking more than 14 cigarettes per day on average.*

#### Supplementary Table S4. Inference results for selected fixed effects or covariates.

| Phenotype | Current smoking status | | Current heavy smoking status | | Global African Ancestry | |
|---|---|---|---|---|---|---|
| | Estimate (S.E.) | P | Estimate (S.E.) | P | Estimate (S.E.) | P |
| $FEV_1$ % predicted | 12.498 (1.371) *** | <2e-16 | 3.975 (0.863) *** | 4.2e-06 | -17.097 (3.467) *** | 8.1e-07 |
| $FEV_1$ | 0.319 (0.038) *** | <2e-16 | 0.123 (0.025) *** | 5.7e-07 | -0.472 (0.099) *** | 1.7e-06 |
| FVC % predicted | 8.191 (1.086) *** | 8.9e-14 | 1.392 (0.726) # | 0.055 | -18.06 (2.882) *** | 4.0e-10 |
| FVC | 0.277 (0.037) *** | 1.5e-13 | 0.066 (0.026) * | 0.011 | -0.602 (0.102) *** | 3.9e-09 |
| $FEV_1$ /FVC | 0.06 (0.008) *** | 5.2e-13 | 0.027 (0.005) *** | 4.3e-09 | -0.007 (0.019) # | 0.68 |

*Estimate: the estimate of effect size; S.E: Standard Error of Estimate; P: p-value of the Likelihood Ratio Test (LRT) when comparing two models, with and without the covariate of interest. Significance codes for p-values: '***' <0.001; '**' <0.01; '*' <0.05; '.' <0.1; '#' not significant.*

#### Supplementary Table S5. Inference results for random effects.

| Phenotype | Heterogeneity 1 | | Heterogeneity 2 | | Heterogeneity | | Medical centers | | Residuals |
|---|---|---|---|---|---|---|---|---|---|
| | Estimate | P | Estimate | P | Estimate | P | Estimate | P | Estimate |
| $FEV_1$ % predicted | 0.38 *** | 2.5e-13 | 0.1 ** | 0.0014 | 0.47 *** | <2e-16 | 0.03 *** | <2e-16 | 0.49 |
| $FEV_1$ | 0.34 *** | 4.9e-10 | 0.05 # | 0.15 | 0.38 *** | 6.7e-14 | 0.03 *** | <2e-16 | 0.59 |
| FVC % predicted | 0.26 *** | 3.4e-06 | 0.1 ** | 0.0099 | - 0.36 *** | 4.0e-11 | 0.03 *** | <2e-16 | 0.61 |
| FVC | 0.22 *** | 0.00016 | 0.02 # | 0.64 | 0.24 *** | 4.9e-05 | 0.03 *** | <2e-16 | 0.73 |
| $FEV_1$ /FVC | 0.53 *** | <2e-16 | 0.12 *** | 2.0e-08 | 0.65 *** | <2e-16 | 0.01 *** | 1.1e-11 | 0.34 |

*The heterogeneity of residual errors among three smoking groups is modeled using three components: the residuals (i.i.d.) (**Residuals** column); extra variance due to current heavy smoking status (**Heterogeneity 2** column); and extra variance due to current smoking status, excluding individuals with heavy smoking status (**Heterogeneity 1** column). Estimate: the estimate of the variance component; P: p-value of the Likelihood Ratio Test (LRT) when comparing two models, with and without the random effect of interest. Significance codes for p-values: '***' <0.001; '**' <0.01; '*' <0.05; '.' <0.1; '#' not significant.*

#### Supplementary Table S6. Fine-mapping (SNP-based) results in loci identified by admixture mapping of ancestry-by-smoking interactions.

| Locus | SNP rank | SNP name | Chr. | Position, bp | Z-score | Posterior Pr. | BF (log10) |
|---|---|---|---|---|---|---|---|
| 1 | 1 | rs933920 | 11 | 12,481,110 | 2.92 | 0.11 | 1.19 |
| 1 | 2 | rs4553350 | 11 | 12,759,834 | -2.84 | 0.09 | 1.08 |
| 1 | 3 | rs6485989 | 11 | 12,695,171 | 2.69 | 0.07 | 0.92 |
| 1 | 4 | rs1564947 | 11 | 12,176,409 | 2.64 | 0.06 | 0.87 |
| 1 | 5 | rs4756884 | 11 | 12,474,098 | -2.55 | 0.05 | 0.78 |
| 2 | 1 | rs7569427 | 2 | 238,413,338 | -2.32 | 0.08 | 0.76 |
| 2 | 2 | rs2280289 | 2 | 238,483,729 | 2.10 | 0.05 | 0.56 |
| 2 | 3 | rs4663794 | 2 | 238,731,931 | -2.03 | 0.05 | 0.49 |
| 2 | 4 | rs4663795 | 2 | 238,733,016 | -1.70 | 0.03 | 0.25 |
| 2 | 5 | rs4663714 | 2 | 238,217,353 | -1.69 | 0.03 | 0.25 |

*The FINEMAP program was used to perform the fine-mapping analysis. Z-score: Z score of association between the SNP-by-smoking interaction term and phenotype; Posterior Pr.: posterior probability that SNP is causal; BF (log10): Bayes Factor at log base 10 scale that supports that SNP is causal. The last two columns are outputs from the FINEMAP program.*

#### Supplementary Table S7. Results on inference of local ancestry by LAMP-LD.

| Ver. | Local Anc. | Anc. Seg. | Long Anc. Seg. | Reference Panel | Genome Build |
|---|---|---|---|---|---|
| 1 | 608,917 | 154,042 | 37,194 | HapMap II (60 CEU, 60 YRI) | hg19 |
| 2 | 675,316 | 229,384 | 30,043 | 1K Genomes III (99 CEU, 108 YRI) | hg19 |

*Two LAMP-LD runs on the COPDGene data were executed, version 1 described in ref.[3] and version 2 presented in this manuscript. The versions differ by the haplotype reference panel used. The inference results in terms of the number of local ancestry segments are similar (all SNPs from the output of LAMP-LD, **Local Anc.** column; all segments from the output of LAMP-LD, **Anc. Seg.** column; long segments (>10,000 bases), **Long Anc. Seg.** column.*

**Supplementary Table S8. Replication of top SNP-smoking interaction signals in the CHARGE consortium[15].**

| Chr. | SNP | SNP *P* | Allele | $Freq_{CEU/YRI}$ | Trait | Exposure | Packs Year; $FEV_1$ | Packs Year; $FEV_1$/FVC |
|---|---|---|---|---|---|---|---|---|
| 11 | rs933920 | *0.0036* | C/T | 0.99/0.89 | $FEV_1$ % predicted | Current smoker | - | - |
| 11 | rs4553350 | *0.0046* | C/T | 0.55/0.95 | $FEV_1$ % predicted | Current smoker | 0.081 | 0.55 |
| 2 | rs7569427 | *0.020* | A/G | 0.94/0.28 | $FEV_1$ | Current heavy smoker | 0.066 | **0.029** |
| 2 | rs2280289 | *0.036* | C/T | 0.84/0.20 | $FEV_1$ | Current heavy smoker | **0.040** | **0.022** |
| 13 | rs1535532 | *0.020* | A/G | 0.67/0.62 | *FVC* | Current heavy smoker | 0.110 | 0.72 |
| 11 | rs11020968 | *0.035* | A/T | 0.84/1.00 | $FEV_1$ | Current heavy smoker | 0.609 | **0.044** |
| 7 | rs10270076 | *0.0046* | A/G | 0.57/0.29 | $FEV_1$ % predicted | Current heavy smoker | 0.424 | 0.43 |
| 8 | rs7000934 | *0.017* | C/T | 0/0.15 | *FVC* | Current heavy smoker | - | - |
| 1 | rs7533237 | *0.0046* | G/T | 0/0.11 | $FEV_1$/FVC | Current smoker | - | - |

*Top SNP-smoking interaction signals for 7 loci identified by admixture mapping. Two top SNPs are listed for the two genome-wide significant loci (the first four rows). The first 7 columns show information of SNP-smoking interactions in the COPDGene study: Chromosome, SNP name, SNP p-value, reference/alternative alleles, reference allele frequencies in two CEU (European) and YRI (African) populations from the 1,000 Genomes Projects, and the trait that drives the signal in the admixture mapping and, thus, was used as outcome phenotype in SNP-smoking interaction association analysis. The last 2 columns show p-values of SNP-smoking interactions in the CHARGE consortium (meta-analysis across 19 studies with the total sample size N = 50,047); the $FEV_1$/FVC outcome phenotype and two smoking exposures are shown. Meta-analysis results are missing for some SNPs, because there SNPs were rare in the European population used in the CHARGE consortium. P-values for SNP-smoking interactions in the CHARGE consortium are marked in bold if they are nominally significant at the level of 0.05.*

#### Table S9. SNP associations reported in GWAS (UK Biobank + SpiroMeta) of either pulmonary phenotypes[12] or COPD[13] and located within 1Mb from the regions detected by admixture mapping in COPDGene.

| Locus | Trait | SNP | Nearest gene | P | Distance (region) | Distance (top ancestry segment) |
|---|---|---|---|---|---|---|
| 11p15.2-3 | COPD | rs4757118 | *ARNTL* | $3.8 \times 10^{-9}$ | 325,400 | 797,555 |
| 11p15.2-3 | COPD | rs80145403* | *PARVA* | $9.8 \times 10^{-6}$ | 0 | 119,611 |
| 11p15.2-3 | COPD | rs7114698* | *TEAD1* | $2.3 \times 10^{-5}$ | 0 | 334,195 |
| 2q37.3 | FVC | rs6431620 | *LINC01107* | $1.4 \times 10^{-10}$ | 83,5077 | 1,118,202 |
| 2q37.3 | $FEV_1$/FVC | rs61332075 | – | $1.7 \times 10^{-9}$ | 546,667 | 829,792 |
| 2q37.3 | COPD | rs72983853* | *ASB1* | $2.5 \times 10^{-5}$ | 554,317 | 837,442 |
| 2q37.3 | COPD | rs13410025* | *TWIST2* | $3.4 \times 10^{-7}$ | 844,365 | 1,127,490 |
| 7p15.2-3 | $FEV_1$ | rs559233 | *SKAP2* | $7.8 \times 10^{-13}$ | 477,550 | 909,106 |
| 7p15.2-3 | FVC | rs62454414 | *HOXA-AS3* | $1.3 \times 10^{-9}$ | 811,049 | 1,242,605 |

***Locus: loci of lung function phenotypes from multi-trait admixture mapping in COPDGene; Trait: phenotype for which GWAS was conducted; SNP: genome-wide significant SNP from GWAS or conditional analysis (marked by *); Nearest gene: the nearest gene to SNP; P: the P-value in GWAS; Distance (region): the distance from SNP to the region (< 1Mb was the criteria to select SNPs); Distance (top ancestry segment): the distance from SNP to the top ancestry segment (multi-trait admixture mapping) in Table 2 (the main text).***

#### Table S10. Marginal effects of local ancestry segments that show top local ancestry-smoking interactions (Table 2 in the main text).

| Locus | Ancestry segment | Trait | Exposure | Z | P |
|---|---|---|---|---|---|
| 11p15.2-3 | 12,075,829-12,845,835 | $FEV_1$ % predicted | Current smoker | 1.55 | 0.12 |
| 2q37.3 | 238,143,387-238,769,892 | $FEV_1$ | Current heavy smoker | -1.66 | 0.09 |
| 13q12.3-13.1 | 31,623,839-32,256,475 | FVC | Current heavy smoker | -0.13 | 0.90 |
| 11q21 | 94,360,812-94,825,729 | $FEV_1$ | Current heavy smoker | 1.46 | 0.14 |
| 7p15.2-3 | 25,133,849-26,371,279 | $FEV_1$ % predicted | Current heavy smoker | 0.24 | 0.81 |
| 8q21.13 | 81,871,222-82,335,354 | FVC | Current heavy smoker | 0.30 | 0.76 |
| 1q44 | 248,020,448-249,208,153 | $FEV_1$/FVC | Current smoker | 0.27 | 0.79 |

***The firsts four columns repeat the information of loci reported in Table 2 in the main text. The last two columns report Z-score and P-value of marginal association between local ancestry segment and outcome phenotype. All interaction- and SNP-related covariates were excluded in the model when performing the marginal association test; the global ancestry covariate was included in the model.***

#### Table S11. Marginal effects of SNPs that show top SNP-smoking interactions (Table 3 in the main text).

| Locus | SNP | Top trait | Exposure | Z | P |
|---|---|---|---|---|---|
| 11p15.2-3 | rs933920 | $FEV_1$ % predicted | Current smoker | 1.48 | 0.14 |
| 11p15.2-3 | rs4553350 | $FEV_1$ % predicted | Current smoker | -0.11 | 0.91 |
| 2q37.3 | rs7569427 | $FEV_1$ | Current heavy smoker | 0.39 | 0.15 |
| 2q37.3 | rs2280289 | $FEV_1$ | Current heavy smoker | 0.79 | 0.43 |
| 13q12.3-13.1 | rs1535532 | FVC | Current heavy smoker | -0.53 | 0.60 |
| 11q21 | rs11020968 | $FEV_1$ | Current heavy smoker | -0.81 | 0.42 |
| 7p15.2-3 | rs10270076 | $FEV_1$ % predicted | Current heavy smoker | 0.15 | 0.88 |
| 8q21.13 | rs10270076 | FVC | Current heavy smoker | -0.28 | 0.78 |
| 1q44 | rs7533237 | $FEV_1$/FVC | Current smoker | -1.59 | 0.11 |

***The firsts four columns repeat the information of SNPs reported in Table 3 in the main text. The last two columns report Z-score and P-value of marginal association between SNP and outcome phenotype. All interaction-related covariates were excluded in the model when performing the marginal association test; both global and correspondent local ancestry covariates were included in the model.***

### Supplementary figures

#### Supplementary Figure S1.

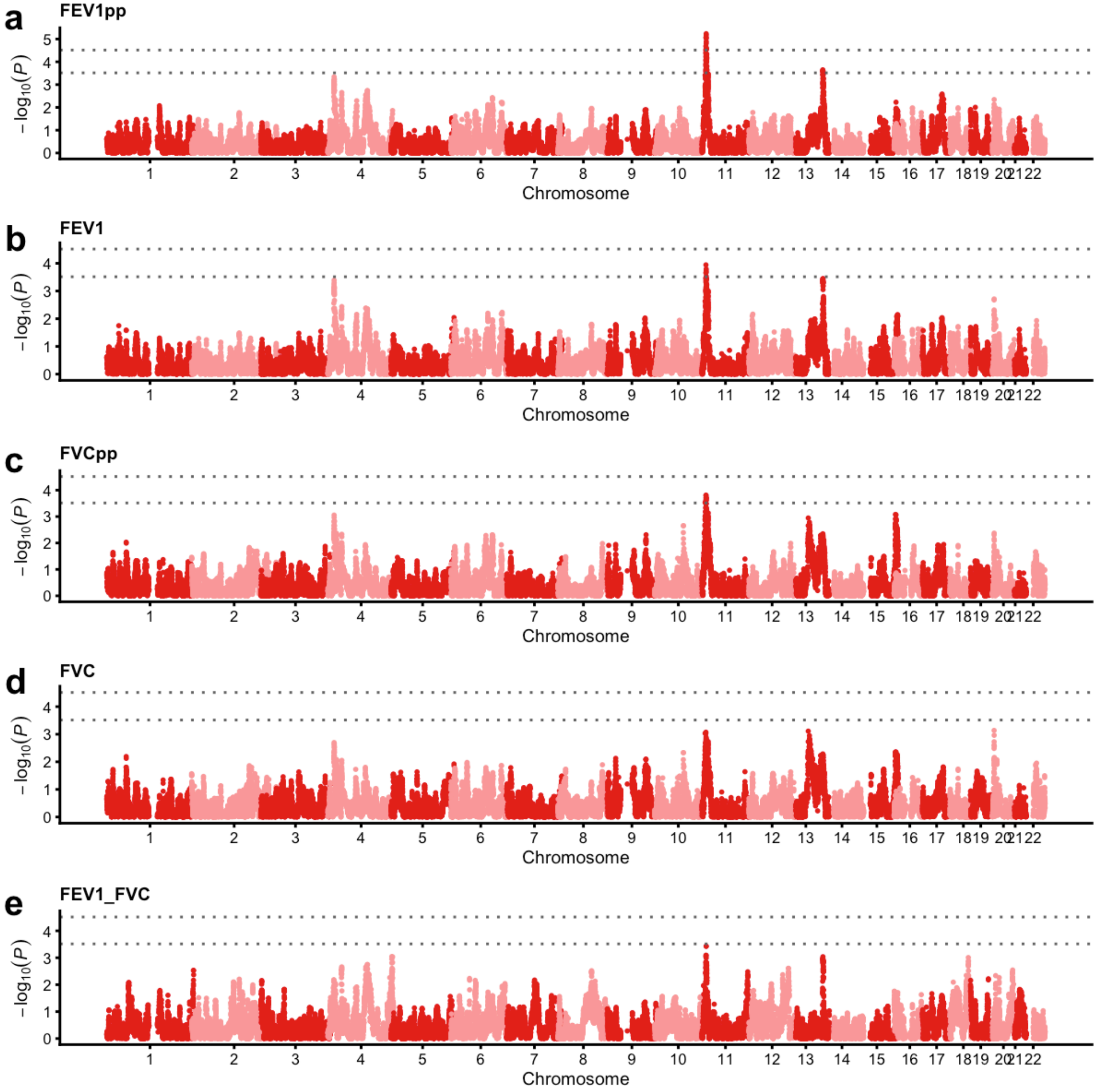


Figure S1: Manhattan plots for single-trait admixture mappings of ancestry-by-smoking interactions, where smoking is the current smoking exposure (current smokers *vs.* current non-smokers). Two horizontal lines depict the genome-wide significant and suggestive Bonferroni thresholds: 0.05/1,635 = 3.06e−5 and 0.5/1,635 = 3.06e−4. The effective number of tests (1,635) is estimated by the eigenMT method.

**Supplementary Figure S2.**

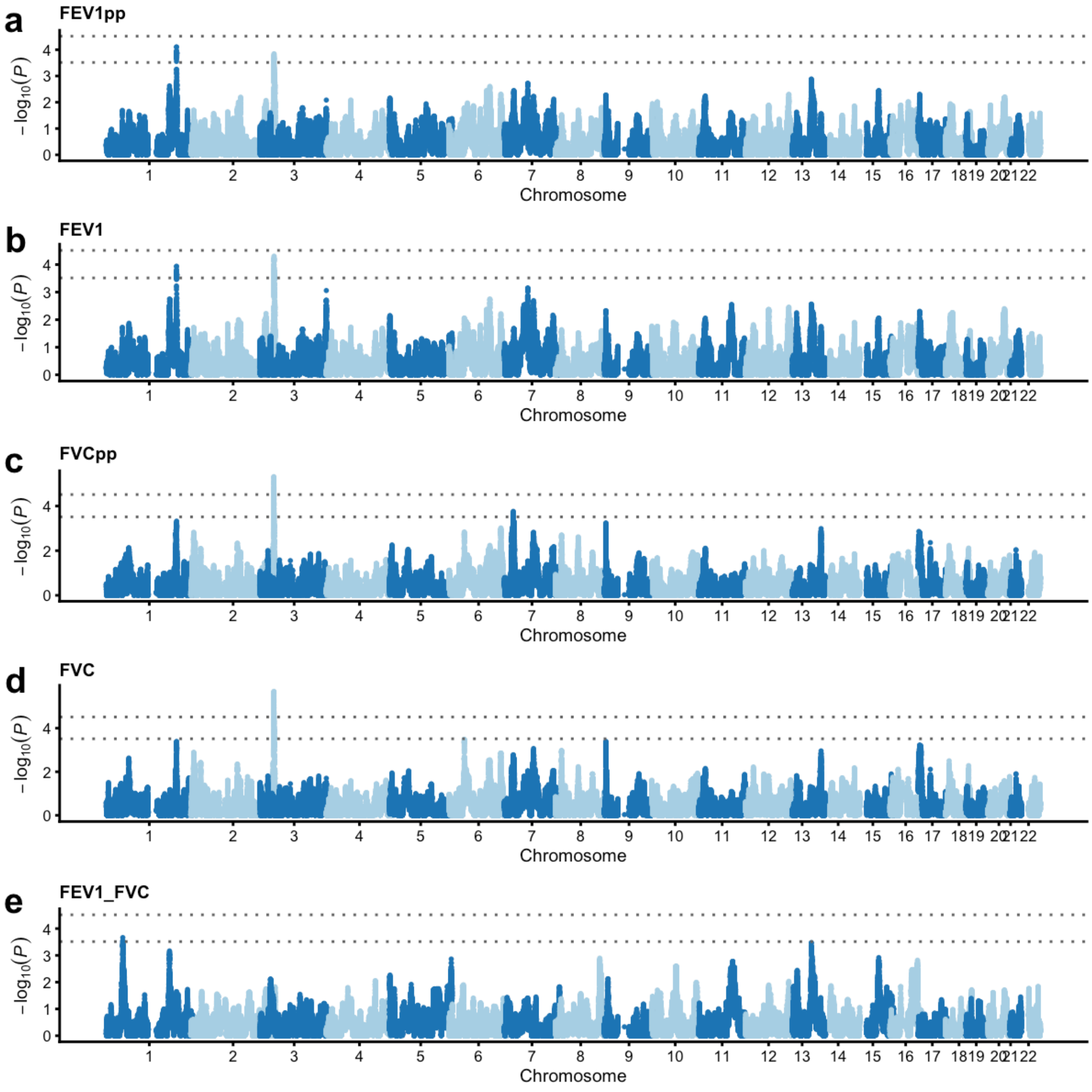


Figure S2: Manhattan plots for single-trait admixture mappings of ancestry-by-smoking interactions, where smoking is the current smoking exposure (current heavy smokers *vs.* current moderate smokers; current non-smokers are ***excluded***). Two horizontal lines depict the genome-wide significant and suggestive Bonferroni thresholds: 0.05/1,635 = 3.06e−5 and 0.5/1,635 = 3.06e−4. The effective number of tests (1,635) is estimated by the eigenMT method.

#### Supplementary Figure S3.

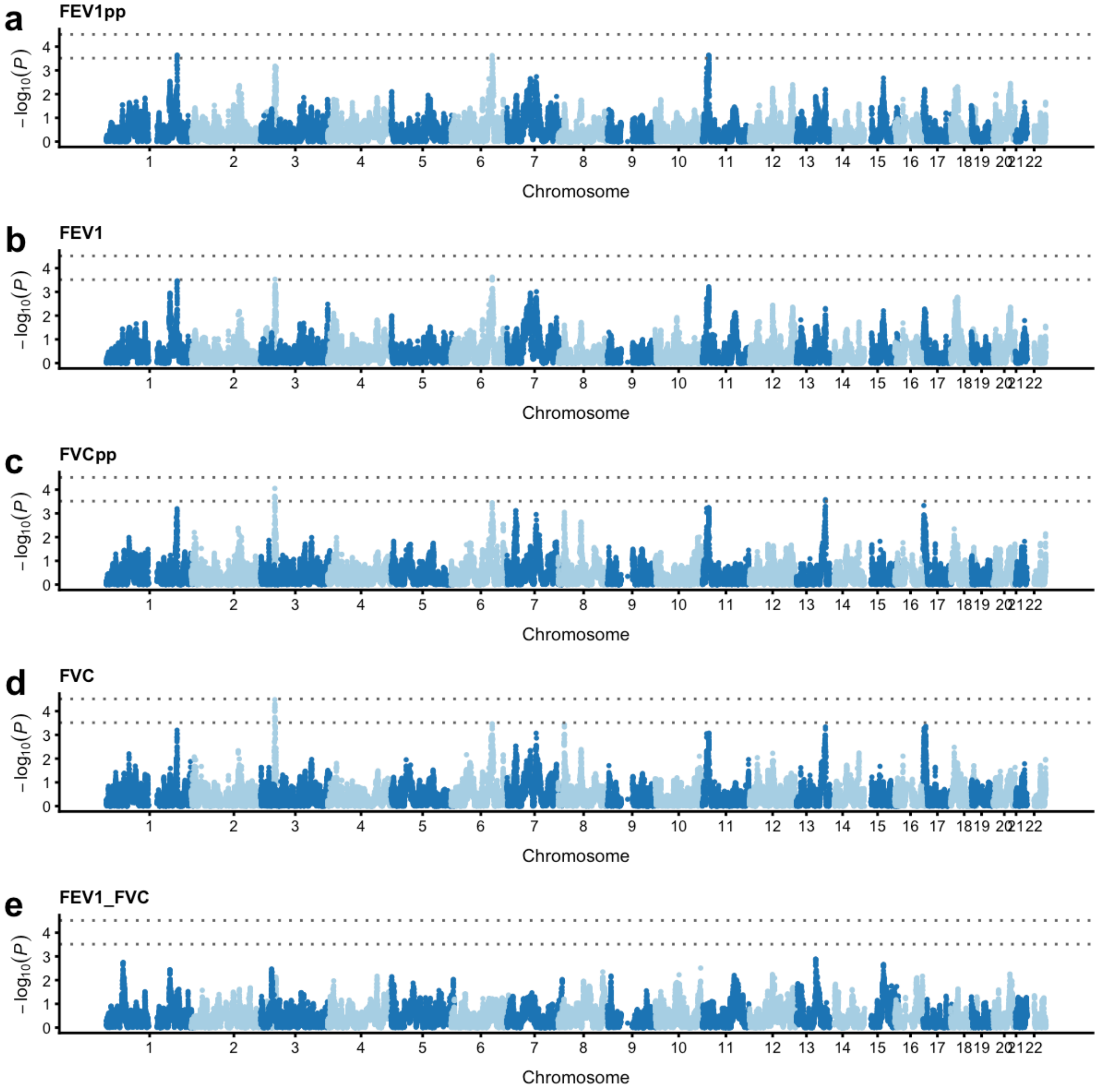


Figure S3: Manhattan plots for single-trait admixture mappings of ancestry-by-smoking interactions, where smoking is the current smoking exposure (current heavy smokers *vs.* current moderate smokers; current non-smokers are ***not excluded***). Two horizontal lines depict the genome-wide significant and suggestive Bonferroni thresholds: 0.05/1,635 = 3.06e−5 and 0.5/1,635 = 3.06e−4. The effective number of tests (1,635) is estimated by the eigenMT method.

**Supplementary Figure S4.**

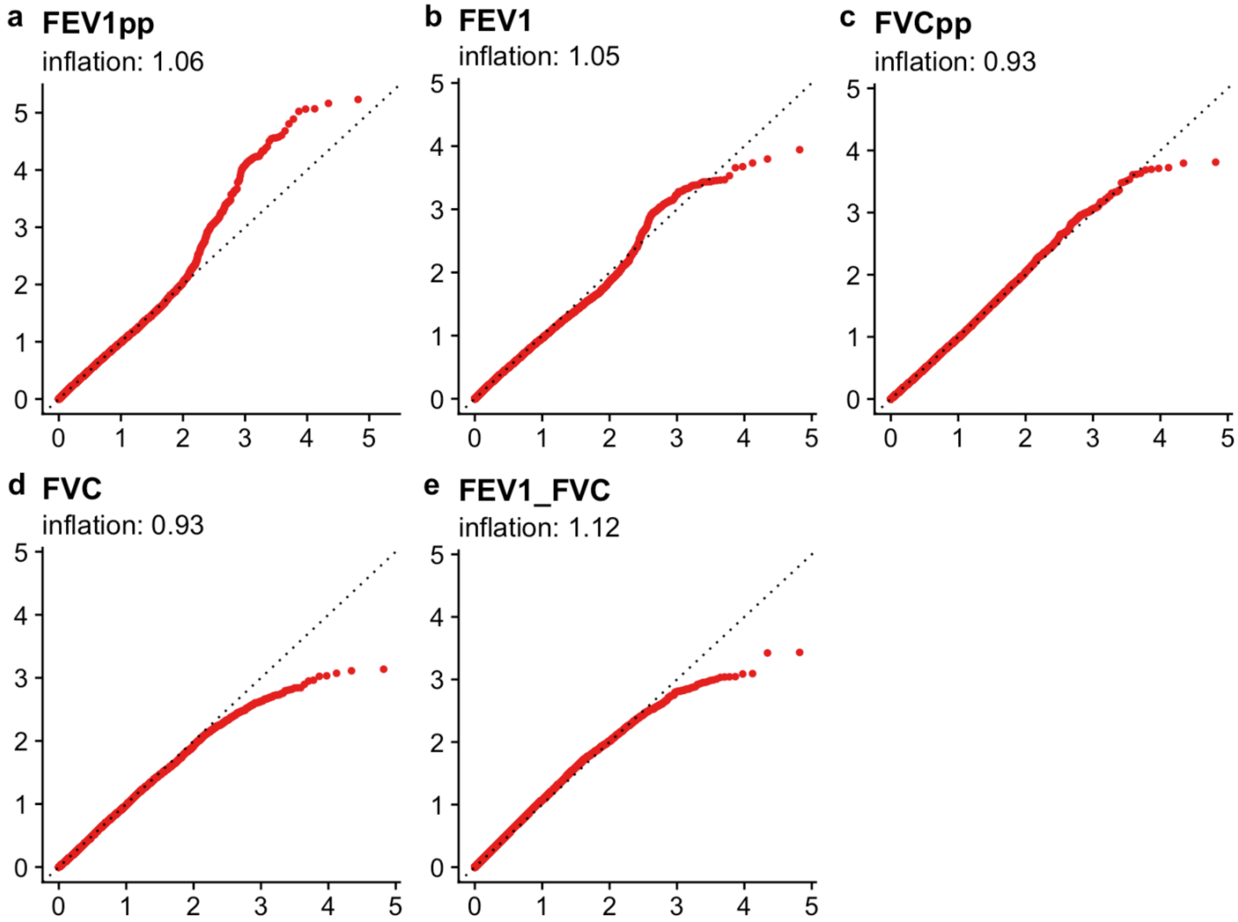


Figure S4: Quantile-quantile (QQ) plots for single-trait admixture mapping of ancestry-by-smoking interactions, where smoking is the current smoking exposure (current smokers *vs.* current non-smokers).

**Supplementary Figure S5.**

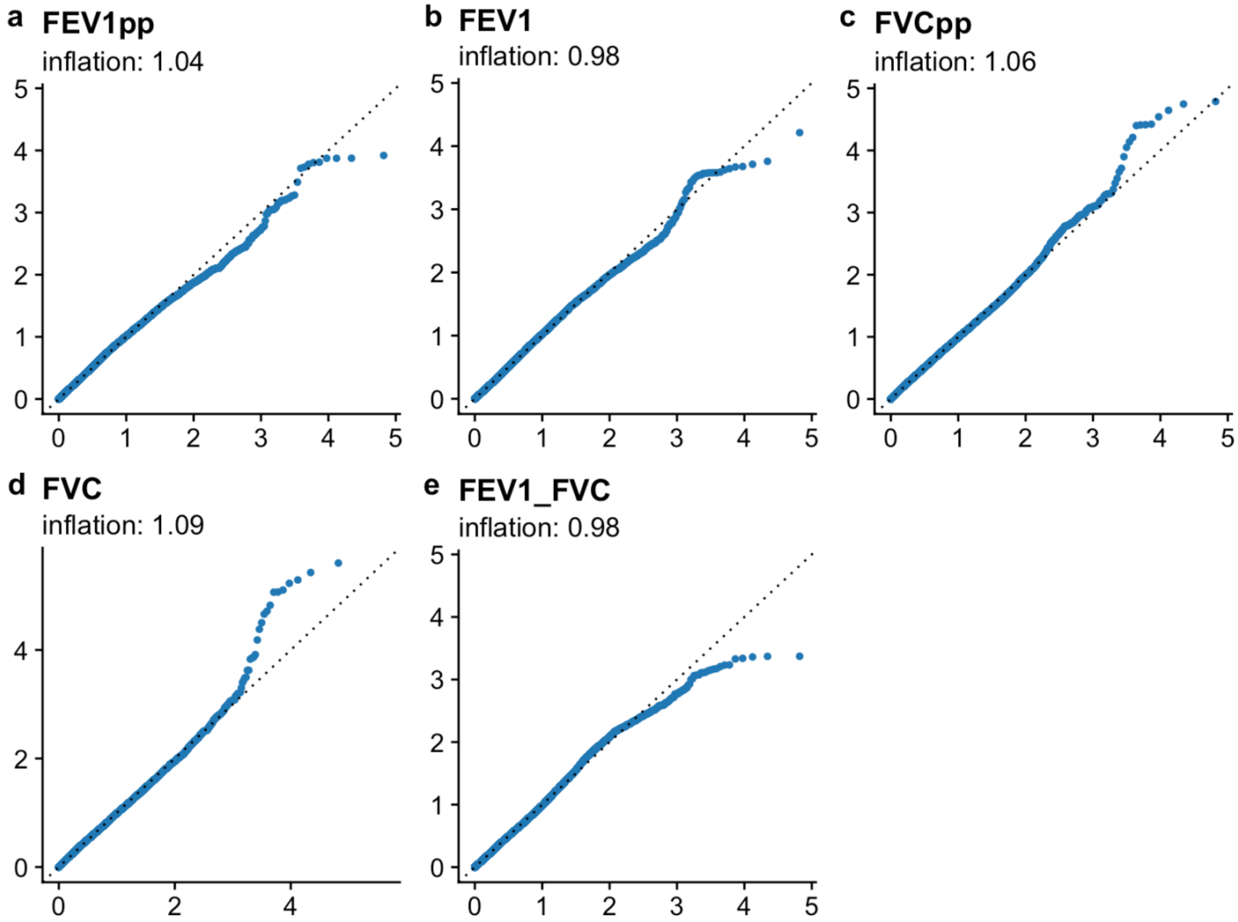


Figure S5: Quantile-quantile (QQ) plots for single-trait admixture mapping of ancestry-by-smoking interactions, where smoking is the current smoking exposure (current heavy smokers *vs.* current moderate smokers; current non-smokers are excluded).

**Supplementary Figure S6.**

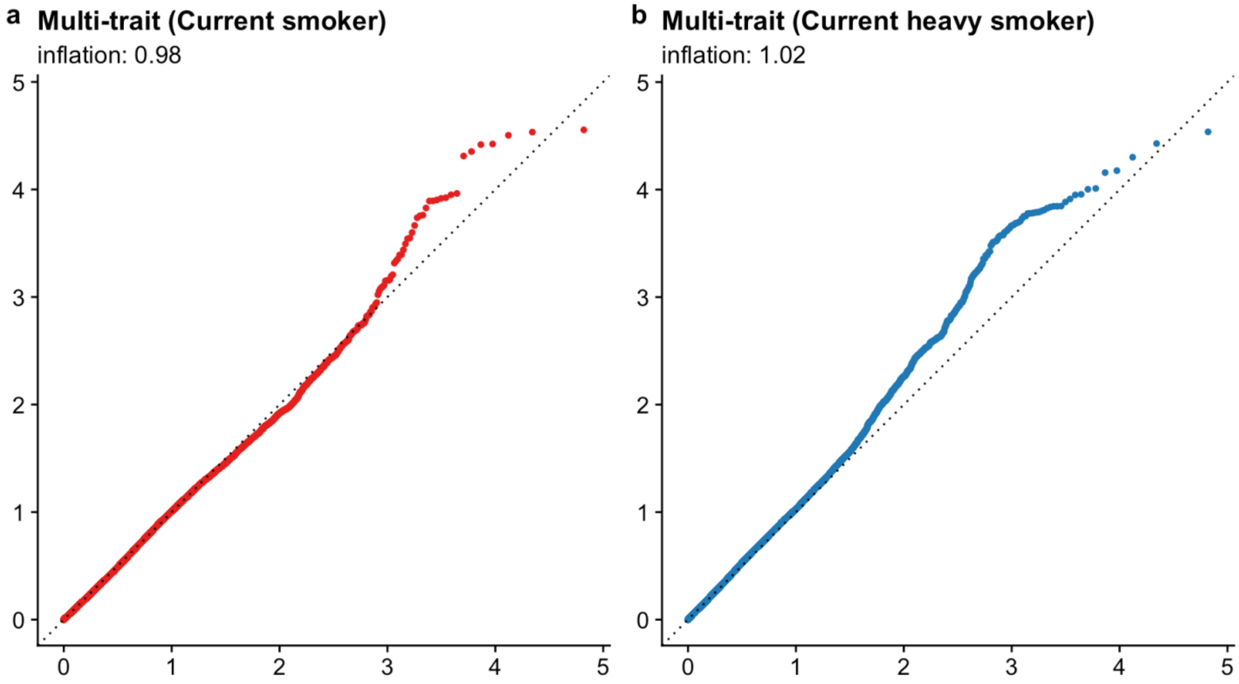


Figure S6: Quantile-quantile (QQ) plots for two multi-trait admixture mappings of ancestry-by-smoking interactions, where smoking is (a) the current smoking exposure (current smokers *vs.* current non-smokers); (b) the current heavy smoking exposure (current heavy smokers *vs.* current moderate smokers; current non-smokers are excluded).

**Supplementary Figure S7.**

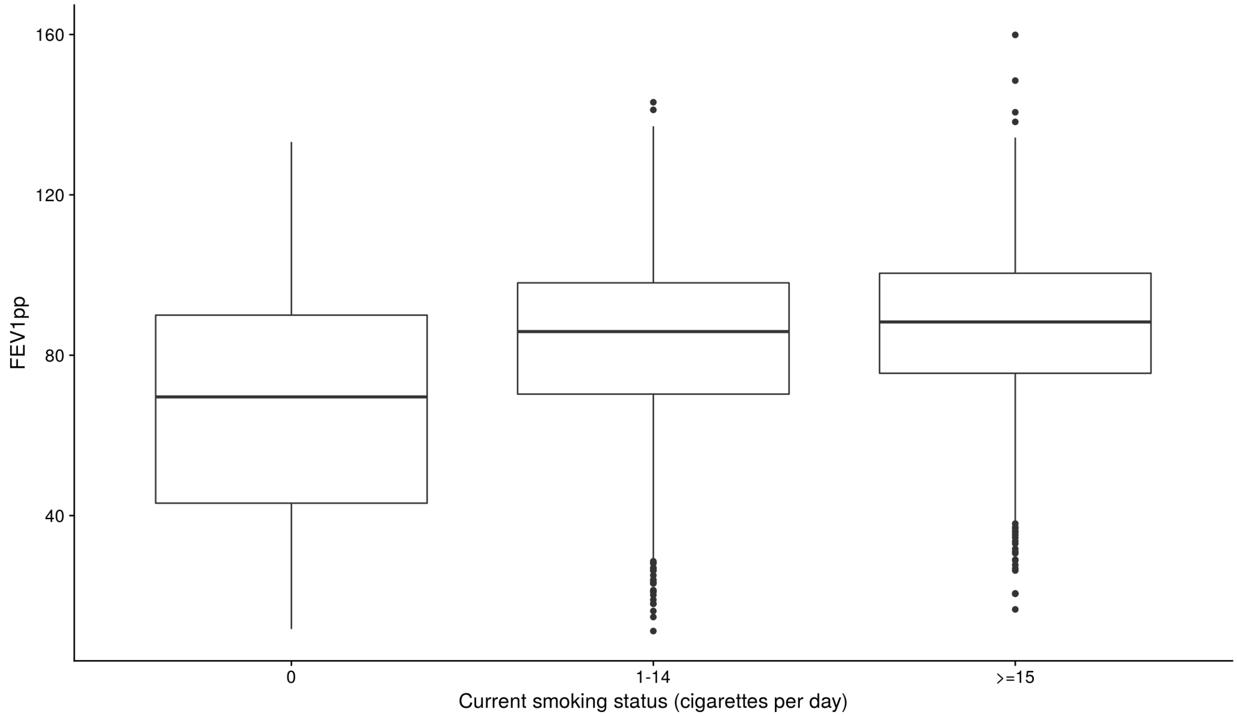


Figure S7: Distribution of FEV1 % predicted trait among three smoking groups: not current smokers; moderate current smokers (1-14 cigarettes per day); and heavy current smokers (>14 cigarettes per day).

**Supplementary Figure S8.**

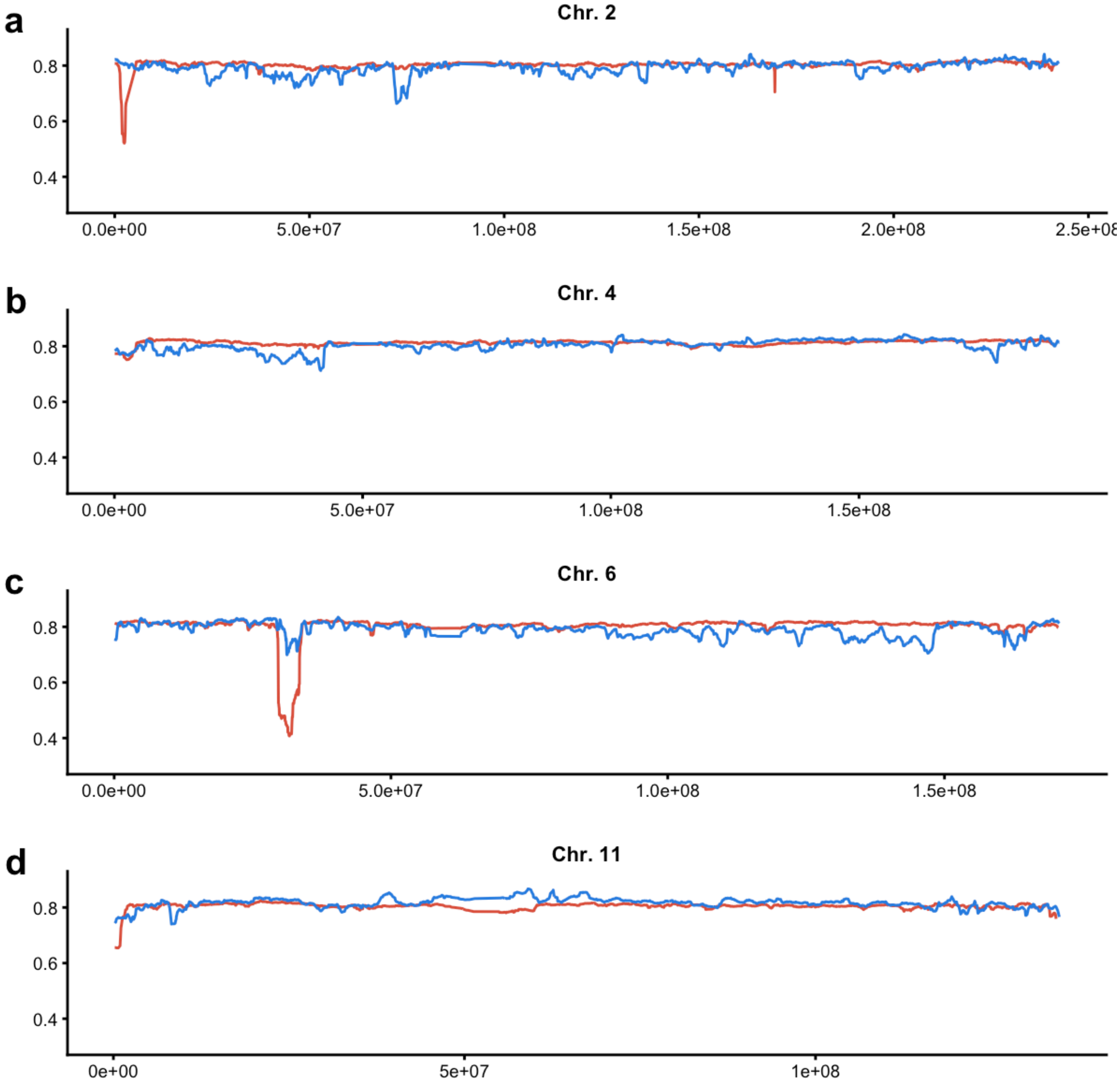


Figure S8: The local ancestry (long) segments averaged across all individuals for Chromosomes (a) 2, (b) 4, (c) 6 and (d) 11. The ancestry inferred using the HapMap II (60 CEU, 60 YRI) is depicted by red lines, and the ancestry inferred using the 1,000 Genomes III (99 CEU, 108 YRI) is depicted by blue lines. See also Supplementary Table S7 for comparison.

**Supplementary Figure S9.**

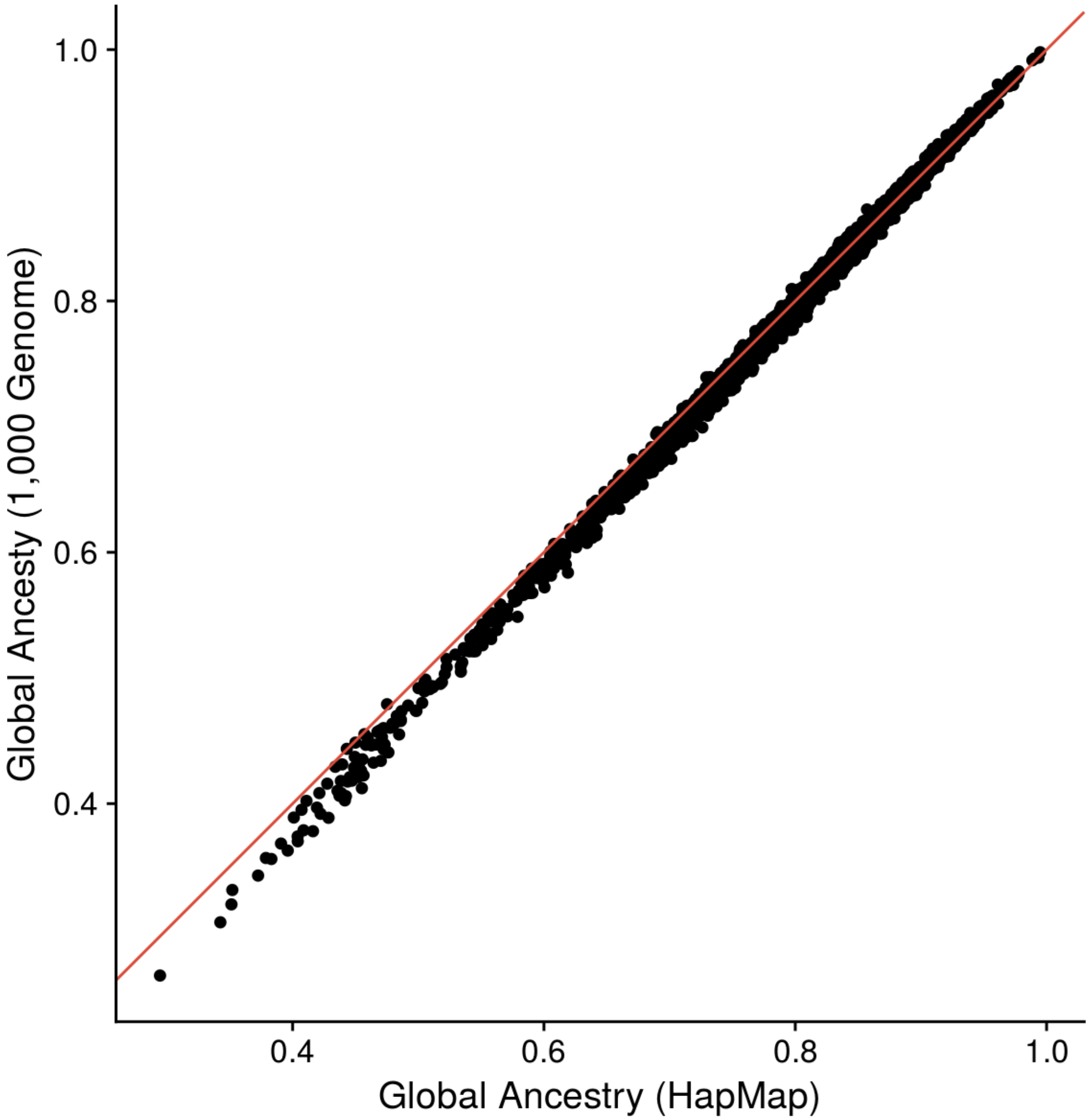


Figure S9: Comparison of the global African ancestry inferred using the HapMap II (60 CEU, 60 YRI) and the 1,000 Genomes III (99 CEU, 108 YRI) reference panels.

#### Supplementary Figure S10.

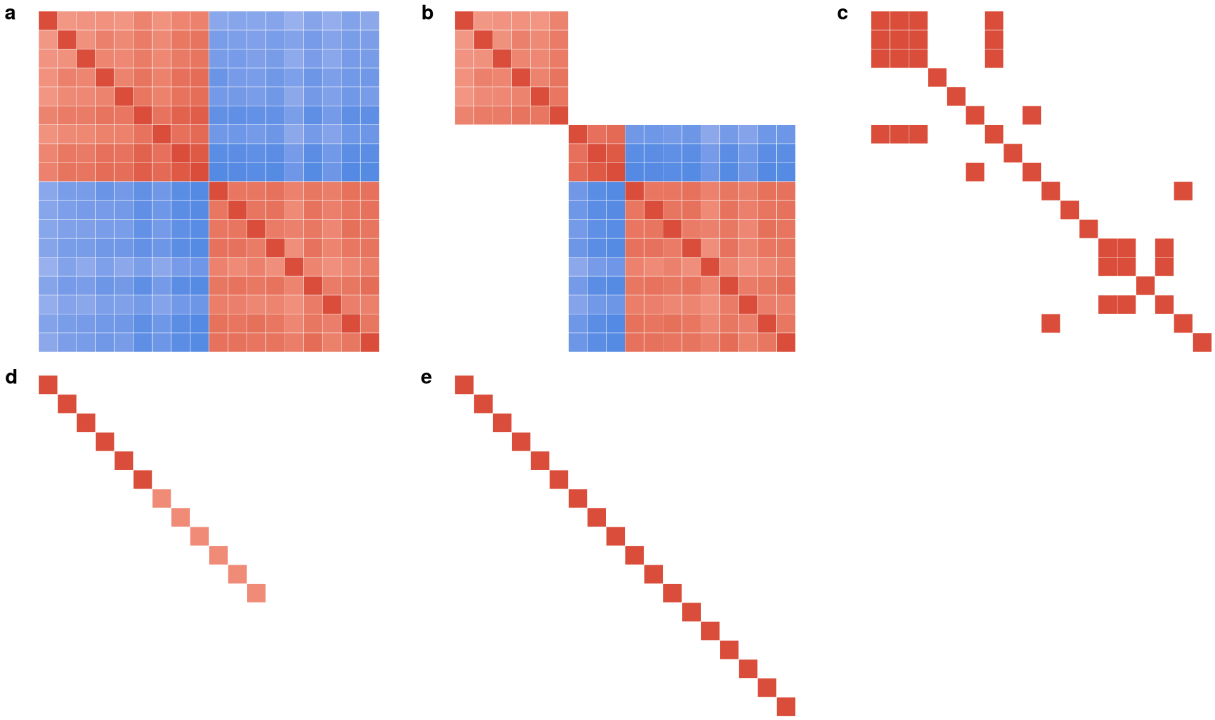


Figure S10: Variance-covariance matrices of the random effects in the linear mixed model (Equation 2 in the main text) developed for admixture mapping of gene-by-environment interactions: (a) the genetic effects with ARM; (b) the genetic effects with EARM; (c) the medical center effects; (d) the heterogeneity effects; and (e) the residual effects. For illustrative purpose only, the relationships are depicted for a subset of 18 individuals, 6 in each of three smoking groups (not current smokers; moderate current smokers with 1-14 cigarettes per day; and heavy current smokers with >14 cigarettes per day). The individuals were selected to enhance the visual contrasts: the first 9 individuals have the largest global African ancestry proportions, while the last 9 individuals have the smallest global African ancestry proportions. Positive and negative values are colored with red and blue, respectively; zero values are left blank.

**Supplementary Figure S11.**

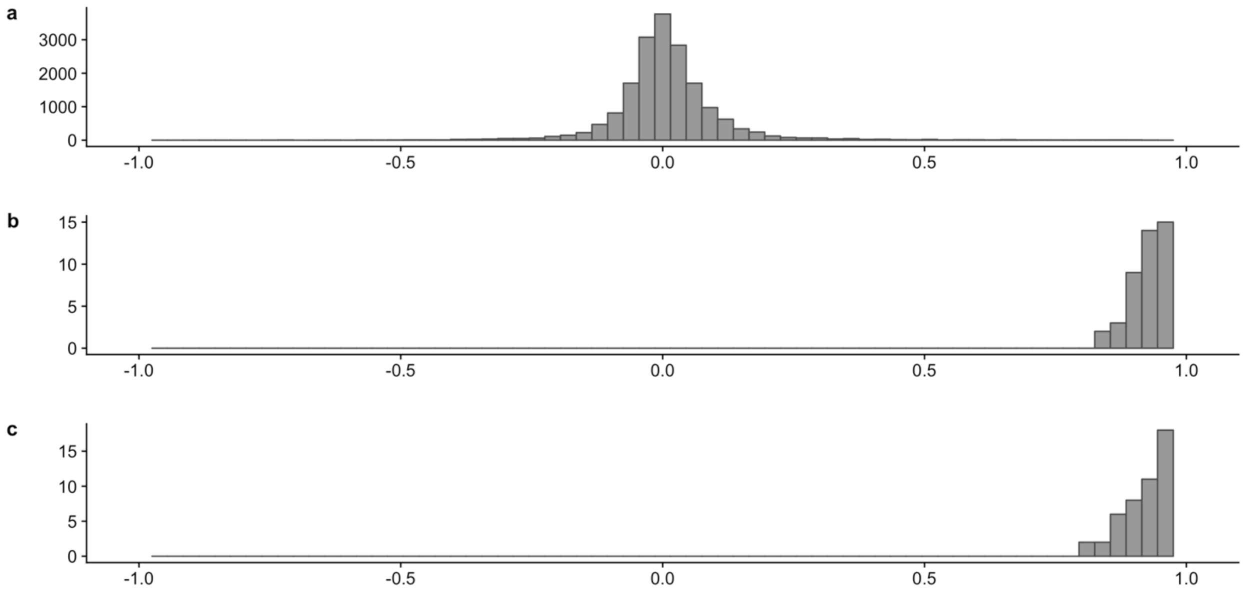


Figure S11: Distribution of correlation among (a) SNPs, (b) local ancestry segments and (c) local ancestry segments corrected for global ancestry (a region of the locus in Chromosome 11:12,075,829-12,845,835).

**Supplementary Figure S12.**

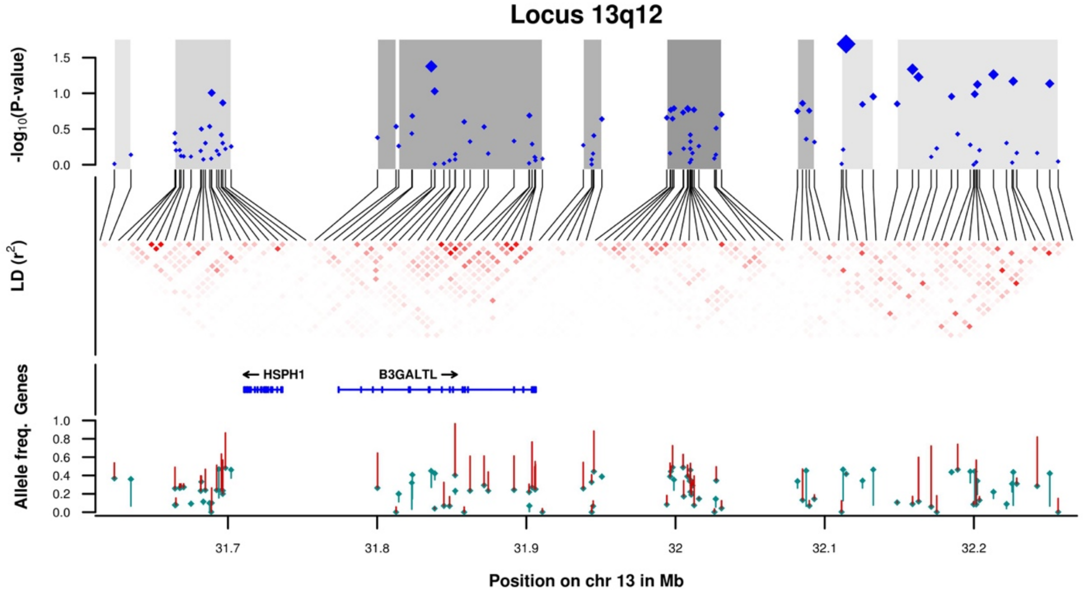


Figure S12. Regional association plot, linkage disequilibrium pattern and trans-continental differences in allelic frequency for the locus 13q12.3-13.1.

**Supplementary Figure S13.**

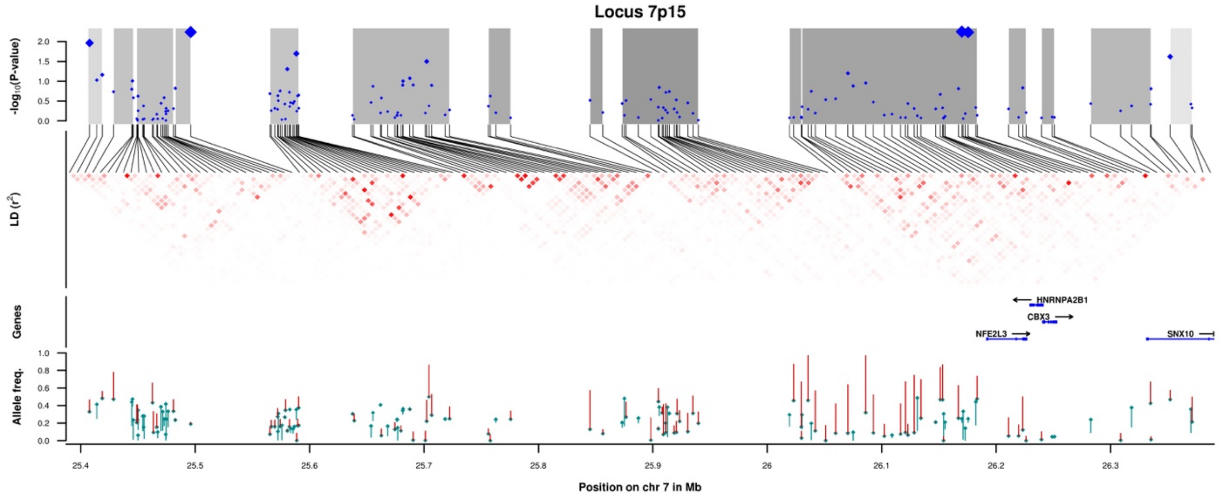


Figure S13. Regional association plot, linkage disequilibrium pattern and trans-continental differences in allelic frequency for the locus 7p15.2-3.

**Supplementary Figure S14.**

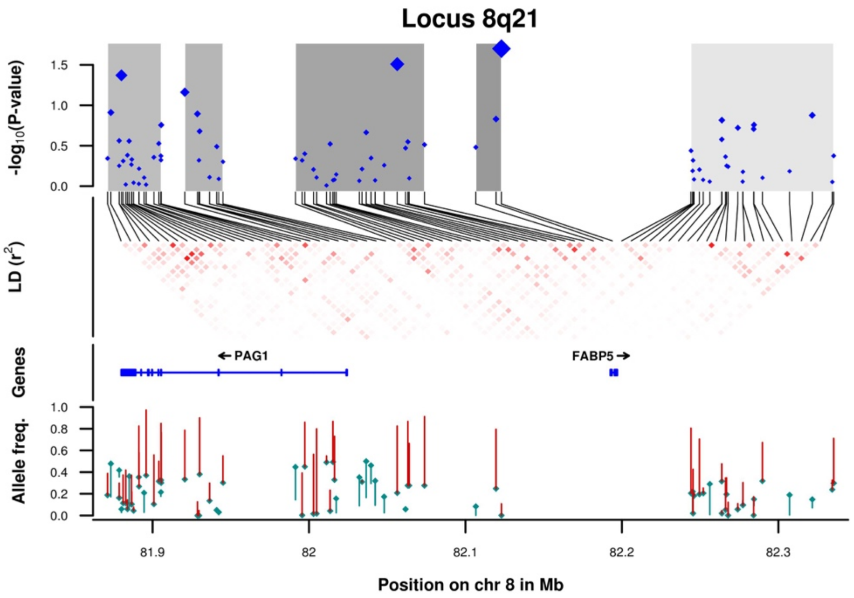


Figure S14. Regional association plot, linkage disequilibrium pattern and trans-continental differences in allelic frequency for the locus 8q21.13.

**Supplementary Figure S15.**

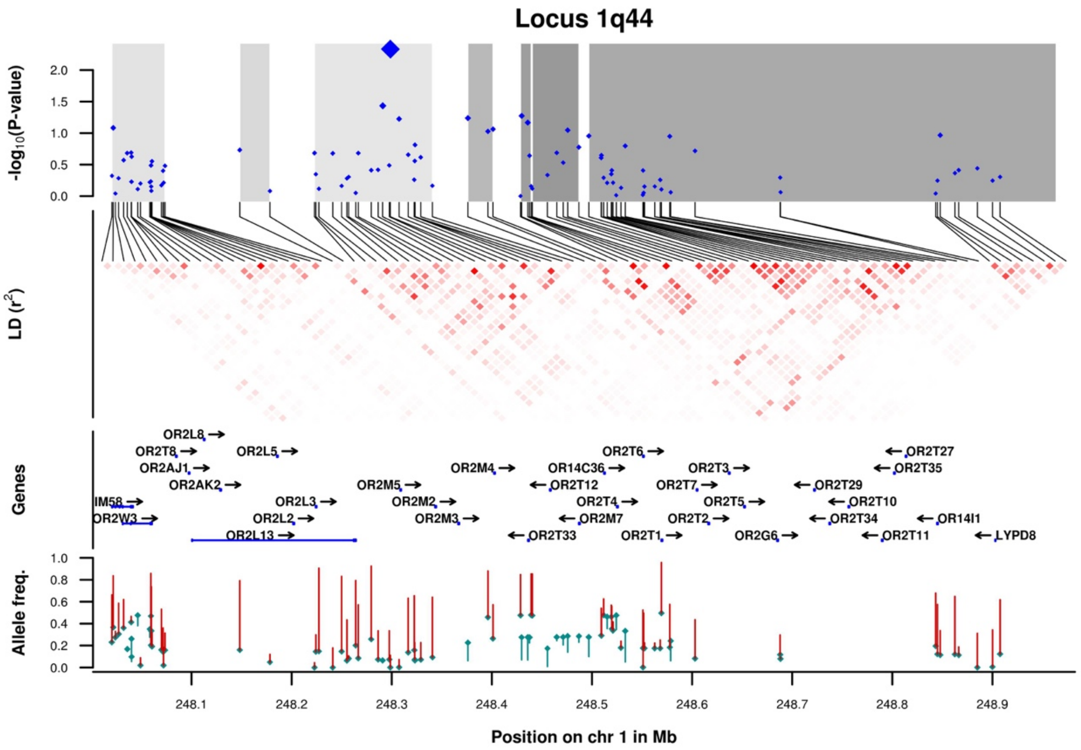


Figure S15. Regional association plot, linkage disequilibrium pattern and trans-continental differences in allelic frequency for the locus 1q44.
